# Ethylene signal-driven plant-multitrophic synergy boosts crop performance

**DOI:** 10.1101/2025.11.28.690471

**Authors:** Marcel Baer, Yanting Zhong, Baogang Yu, Tian Tian, Xiaoming He, Ling Gu, Xiaofang Huang, Elena Gallina, Isabelle Metzen, Marcel Bucher, Rentao Song, Caroline Gutjahr, Zhen Su, Yudelsy A.T. Moya, Nicolaus von Wirén, Siji Wang, Zheng Zhao, Lin Zhang, Lixing Yuan, Yunlu Shi, Mareike Baer, Weiwei Qi, Chunjian Li, Xuexian Li, Frank Hochholdinger, Peng Yu

## Abstract

Efficient nutrient use in agriculture depends on the dynamic interplay between plant roots, soil, and microbial communities. The root–rhizosphere interface is central to nutrient uptake and serves as a key hub for interactions with beneficial microbes. Arbuscular mycorrhizal (AM) fungi, positioned at the nexus between plant roots and soil microbiota, play a critical role in enhancing crop performance under nutrient-limited conditions. In this study, we dissected the genetic and molecular basis of AM fungi–induced lateral root development in maize (*Zea mays*), focusing on the role of ethylene-responsive transcription factors (ERFs). We identified ERF genes as essential regulators of pericycle cell division, acting downstream of ethylene biosynthesis genes (*ACS6* and *ACS7*) and AM fungal signaling. Our findings reveal that AM fungi promote lateral root initiation by activating ERF expression and reprogramming flavonoid metabolism, particularly reducing the accumulation of flavonols such as kaempferol and quercetin, which otherwise inhibit root development when over-accumulated. Furthermore, we demonstrated that *Massilia*, a beneficial rhizobacterium, synergizes with AM fungi to enhance lateral root formation by colonizing fungal hyphae, degrading flavonoids, and contributing to auxin production. Together, our results uncover a tripartite signaling network linking ethylene signaling, flavonoid-mediated microbial recruitment, and AM symbiosis. This study highlights ERFs as central integrators of plant–microbe interactions and provides a molecular framework for engineering root architecture and microbiome assembly to improve nutrient acquisition and support sustainable crop productivity.

## Introduction

Arbuscular mycorrhizal fungi (AMF) form ancient and widespread mutualistic associations with the majority of land plants, enhancing nutrient acquisition, especially phosphorus, and contributing to plant stress resilience (Lanfranco et al., 2018). Through co-evolution and domestication, AMF have specialized as obligate biotrophs that rely on plant roots for carbon while reciprocally supporting host plants with water and mineral nutrients (Gutjahr and Parniske, 2013). Beyond this well-established plant–fungus symbiosis, AMF also associate with diverse soil bacteria, forming tripartite interactions that further enhance nutrient cycling and plant performance (Duan et al., 2024). These mycorrhiza-associated bacteria, often colonizing the hyphosphere or forming biofilms on extraradical hyphae, contribute to phosphorus solubilization (Sun et al., 2025), microbial community structuring (Desirò et al., 2014; Xue et al., 2019), and plant immunity (Zhang et al., 2025). Thus, AMF function as a central biological hub that connects root development with beneficial microbial networks, playing a critical role in sustainable crop productivity (Genre et al., 2020).

The spatial complexity of cereal root systems is essential not only for optimizing soil resource capture (Hochholdinger et al., 2018), but also for supporting the establishment of functional mycorrhizal symbioses (Yu et al., 2016). However, the molecular mechanisms governing AMF-mediated root system remodeling, especially in non-legume crops like maize, remain poorly understood. The initial stages of mycorrhizal establishment involve strigolactones released by the plant and fungal signals such as chitooligomers (COs) and lipochitooligosaccharides (LCOs) (MacLean et al., 2017). These early signals trigger cellular changes in both partners, but how this leads to reprogramming of root developmental trajectories is not fully resolved.

In this context, flavonoids have emerged as critical plant-derived metabolites that influence AMF development, bacterial recruitment, and root–rhizosphere communication (Hassan and Mathesius, 2012; Tian et al., 2021; Cook et al., 2025). Beyond their classical role in legume–rhizobia interactions, flavonoids also promote AMF spore germination and hyphal branching, while shaping rhizosphere bacterial communities, including beneficial genera such as *Massilia* and *Pseudomonas* (Yu et al., 2021; He et al., 2024; Zhao et al., 2024; Wu et al., 2025). These findings suggest that flavonoids act as integrative molecular hubs, coordinating plant–fungus–bacterium interactions that underlie adaptive root responses.

Lateral root development, which enhances root system plasticity and nutrient foraging, is especially sensitive to environmental and microbial cues. In maize, lateral roots are initiated from phloem pole pericycle cells and are regulated by both auxin-dependent and auxin-independent pathways (Yu et al., 2016; Van Norman et al., 2013). The maize *lateralrootless1* (*lrt1*) mutant lacks embryonic lateral roots, yet shows partial restoration of lateral root formation upon AMF inoculation, even under low-phosphorus conditions (Paszkowski and Boller, 2002). Interestingly, this restoration does not involve auxin supplementation or altered PIN localization, suggesting that alternative host signaling mechanisms drive AM-induced lateral root formation in *lrt1* (Hochholdinger and Feix, 1998; Schlicht et al., 2013).

In this work, we report for the first time that members of the host *ERF* (ethylene response factor) gene family serve as key regulators of interkingdom communication between maize roots, AMF, and rhizosphere bacteria. Through a combination of genetic, transcriptomic, metabolomic, and microbial profiling experiments, we demonstrate that *ERF* transcription factors integrate flavonoid-mediated microbial recruitment, ethylene signaling, and root developmental reprogramming during AM symbiosis. Our data reveal that AMF remodel root developmental programs by reshaping flavonoid homeostasis, which in turn recruits *Massilia* spp. that form biofilms on fungal hyphae, promote auxin production, and enhance phosphorus solubilization in the rhizosphere. This study uncovers a tripartite signaling framework in which flavonoids act as central mediators between AM fungi, beneficial bacteria, and host developmental genes. Notably, we identified *ERF97* and *ERF147* as downstream targets of ethylene biosynthesis genes (*ACS6/ACS7*), forming a regulatory cascade that links environmental nutrient status to root architectural adaptation. These findings highlight a previously unknown function of host *ERF* genes in coordinating mycorrhizal and bacterial interactions, thus marking a significant advance in our understanding of plant–microbiome communication in cereals. By elucidating how plant signaling pathways and secondary metabolites orchestrate multi-kingdom symbioses, our work offers new molecular targets for engineering root traits and optimizing microbiome-assisted crop performance, particularly under nutrient-limited conditions.

## Results

### Pericycle division competence is coupled with stage-specific transcriptomic reprogramming during lateral root initiation

To elucidate the molecular basis of lateral root initiation in maize, we analyzed the spatiotemporal pattern of pericycle cell division in the lateral root-deficient mutant *lrt1* compared to its wild type (WT). Histological examination revealed that in *lrt1*, pericycle cell divisions are consistently arrested at stage II, along the entire longitudinal developmental zone of the primary root axis, leading to a failure in lateral root primordium formation (Fig. 1A). To explore the transcriptional basis of this defect, we performed laser capture microdissection (LCM) of phloem-pole (PP) and xylem-pole (XP) pericycle cells at defined lateral root initiation stages (5–10 mm, 10–15 mm, and 15–20 mm from the root tip, corresponding to stages I–III; Fig. 1B), followed by RNA-seq. Principal component analysis (PCA) demonstrated that pericycle cell identity (PP vs. XP) accounted for the largest variance (PC1, 29.8%), while genotype-dependent differences (*lrt1* vs. WT) explained additional variance (PC2, 9%) (Fig. 1C). Notably, transcriptome profiles of *lrt1* PP pericycle cells appeared stalled during the stage II–III transition, whereas WT PP pericycle cells exhibited clear transcriptional progression corresponding to lateral root initiation. These findings suggest that in maize, the transcriptomic reprogramming of pericycle cells is tightly linked to their competence to divide, and that *lrt1* disrupts this coordination, blocking the developmental progression required for lateral root formation.

**Figure 1.**
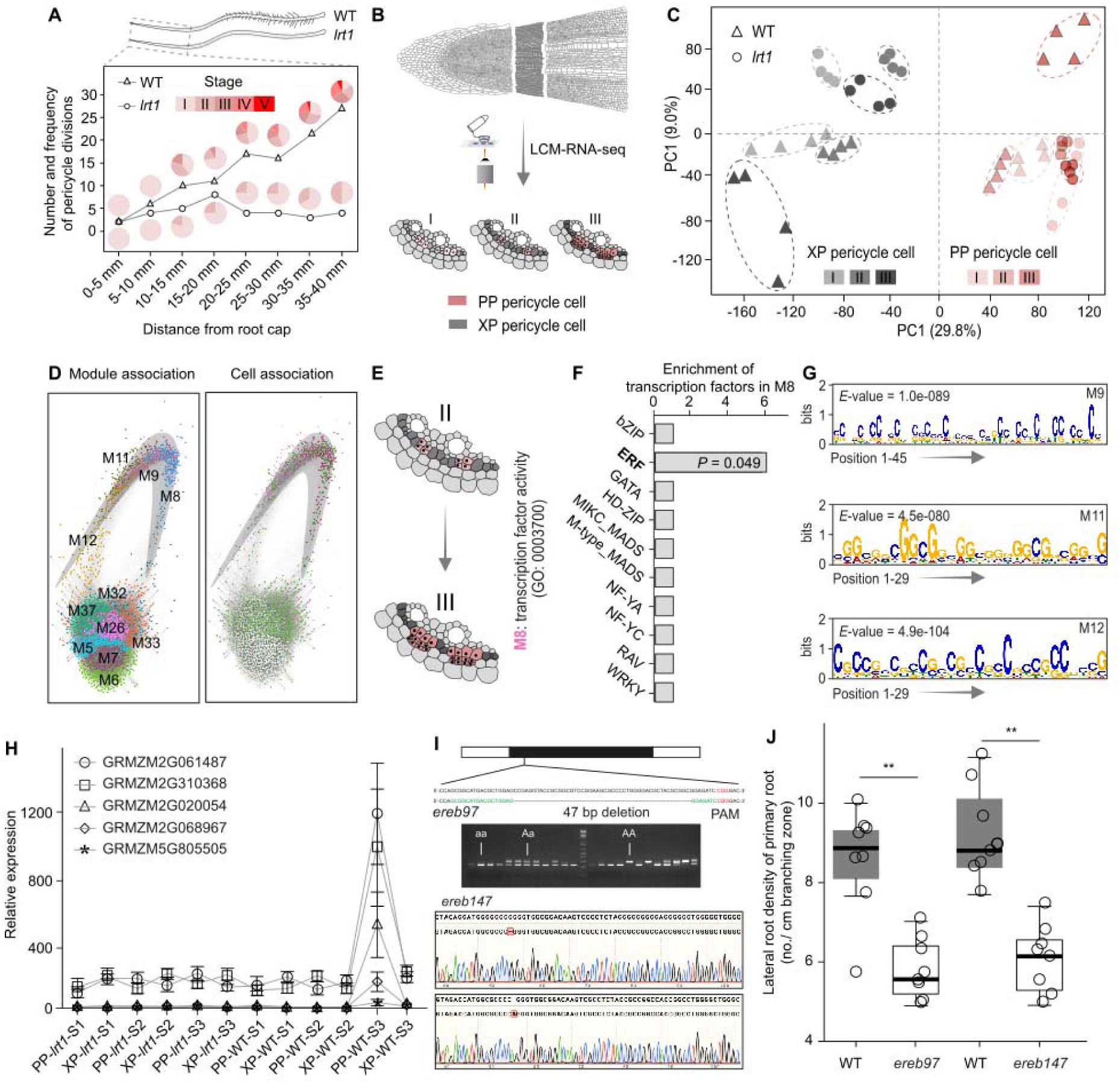
Ethylene response factors – mediated transcriptional regulation of lateral root initiation in maize. (**A**) Developmental stages of lateral root formation in the maize *lrt1* mutant and wildtype. Relative proportion of developmental stages of lateral root primordia observed in 10-day-old maize seedlings of the wild-type B73 and the mutant *lrt1*. Primordia were examined and counted from paraffin sections using at least 6 plants for each genotype. Developmental stages nomenclature of maize lateral root was according to Yu et al., 2016. (**B**) RNA-seq analysis of stage-dependent phloem pole (PP) and xylem pole (XP) pericycle cells isolated by laser capture microdissection (LCM). Pericycle cell types were captured from the same cryo-sections to have highly accurate comparison. The detailed procedure of LCM was described in the Methods. (**C**) Principal component analysis (PCA) for pericycle cell transcriptomes from different developmental stages of maize *lrt1* mutant and wildtype. *n* = 4 biologically independent samples collected from at least 25 continuous cryo-sections per sample. (**D**) Gene module network and cell trait associations by weighted gene co-expression network analysis (WGCNA) and Spearman correlation coefficient (SCC) analysis. Different gene modules were indicated by different colour codes. Module eigengene expression was correlated with lateral root initiation status of *lrt1* mutant and wildtype and clustered from green (SCC = 0) to pink (SCC = 1). (**E**) Identification and characterization of gene module 8 in association with pericycle cell developmental transition from stage II to stage III. Significantly (FDR adjusted *P* = 0.047) enriched gene ontology (GO) term is annotated into “transcription factor activity” (GO:0003700). (**F**) Statistically enrichment analysis of different transcription factor families in module 8 based on the maize transcription factor background (https://planttfdb.gao-lab.org/). Significant *P* value was corrected by FDR correction. ERF, ethylene response factor. (**G**) Gene motif enrichment analysis for M8 coregulated modules (M9, M11 and M12). Gene promoter region was extracted and subjected to MEME for detecting significantly enriched putative binding motif by CentriMo. Significant enrichments were measured by *E*-value applying Bonferroni correction to the term’s *P*-value. (**H**) Relative expression of ERF genes during lateral root development in maize *lrt1* mutant and wildtype. PP, phloem pole; XP, xylem pole; S, stage. (**I**) Production of ERF mutants via CRISPR/Cas9. PAM, protospacer-adjacent motif. (**J**) Lateral root density phenotype of *ereb97* and *ereb137* mutants. n = 8. Statistical significance was controlled by paired Student’s *t* test.

### *ERF* transcription factors orchestrate pericycle cell transition during lateral root initiation

To identify transcriptional regulators associated with *LRT1*-dependent pericycle competence, we applied weighted gene co-expression network analysis (WGCNA) to the pericycle cell transcriptomes. This analysis revealed 39 distinct co-expression modules representing coordinated gene expression programs across developmental stages and genotypes (Fig. 1D). Among these, modules M5–M7 were preferentially enriched in dividing phloem-pole (PP) pericycle cells, whereas M26, M32, M33, and M37 were associated with xylem-pole (XP) pericycle identity (Supplemental Fig. 1). Notably, the modules M8, M9, M11, and M12 were specifically activated during stage III of PP pericycle cells in wild-type roots (Fig. 1D), coinciding with the onset of lateral root stem cell niche establishment (Fig. 1A), suggesting a functional role in driving cell division competence. Gene Ontology (GO) analysis of stage III-enriched modules revealed significant enrichment for biological processes including cell cycle progression (M5), microtubule cytoskeleton organization (M6), and regulation of mitotic transitions (M7) (Supplemental Table 1), supporting the cell division status of these co-expression modules from PP pericycle cells. In contrast, XP-enriched modules were associated with cell wall biogenesis (M26) and hormone/auxin signaling pathways (M32, M33), highlighting the cell-type specificity of transcriptional programs (Supplemental Table 1).

Interestingly, promoter motif enrichment analysis of the three PP-specific modules (M9, M11, M12) identified a conserved GCC-rich cis-regulatory motif, which closely resembles known binding sites of ethylene response factors (ERFs) (Fig. 1G; Supplemental Table 2). Further analysis revealed that Module 8, co-regulated with these modules, was significantly enriched for genes annotated with “transcription factor activity” (GO:0003700) (Fig. 1E), with ERF genes representing the most significantly overrepresented transcription factor family (Fig. 1F; Supplemental Table 3). Among these, five ERF genes exhibited strong expression peaks during stage III of PP pericycle cells in wild-type plants (Fig. 1H), precisely at the developmental window where lateral root initiation occurs, implicating them as potential regulators of pericycle cell division.

To functionally validate the general role of ERF genes in lateral root development, we performed tissue-specific RNA-seq from dissected stele tissues of maize representative IBM recombinant inbred lines (RILs) exhibiting either high or low lateral root density (Supplemental Fig. 2a-c). Comparative transcriptomic analysis identified over 1,100 differentially expressed genes, among which genes encoding ERF and MYB transcription factors were significantly enriched in high lateral root density genotypes (FDR = 0.03, Fisher’s exact test < 0.05) (Supplemental Fig. 2d-g), suggesting ERF-mediated transcriptional control may represent a conserved mechanism underlying lateral root initiation in maize. Finally, we generated CRISPR/Cas9 knockout lines for two ERF genes, *ereb97* and *ereb147*, both highly expressed in competent pericycle cells (Fig. 1I). Both mutants showed a significant reduction (~30%) in lateral root density on the primary root compared to wild-type controls (Fig. 1J), confirming a positive regulatory role of ERF genes in lateral root formation. Together, these findings establish ERFs as key transcriptional regulators linking cell-specific gene expression dynamics to the developmental progression of pericycle cells, thereby promoting lateral root initiation in maize.

### AM fungi partially restore lateral root formation through ERF activation

To further investigate the role of AM fungi in this recovery process of lateral root development, we inoculated *lrt1* mutant plants with AM fungi in phosphate-deficient artificial soil. Six-week-old *lrt1* plants displayed significantly increased lateral root formation (Fig. 2A) and enhanced shoot biomass upon AM fungal inoculation (Supplemental Fig. 3), indicating that AM symbiosis can partially rescue root developmental defects and promote growth under low-phosphorus conditions. To uncover the molecular underpinnings of this interaction, we again used laser capture microdissection (LCM) to isolate pericycle cells at developmental stages I–III from both wild-type and *lrt1* roots, followed by transcriptome profiling via RNA-seq. Principal component analysis revealed that AM fungal inoculation imposed a stronger transcriptional shift on pericycle cells than the genotypic differences between *lrt1* and wild type (Fig. 2B), suggesting that AM fungi exert a dominant influence on pericycle cell transcriptomic programming.

**Figure 2.**
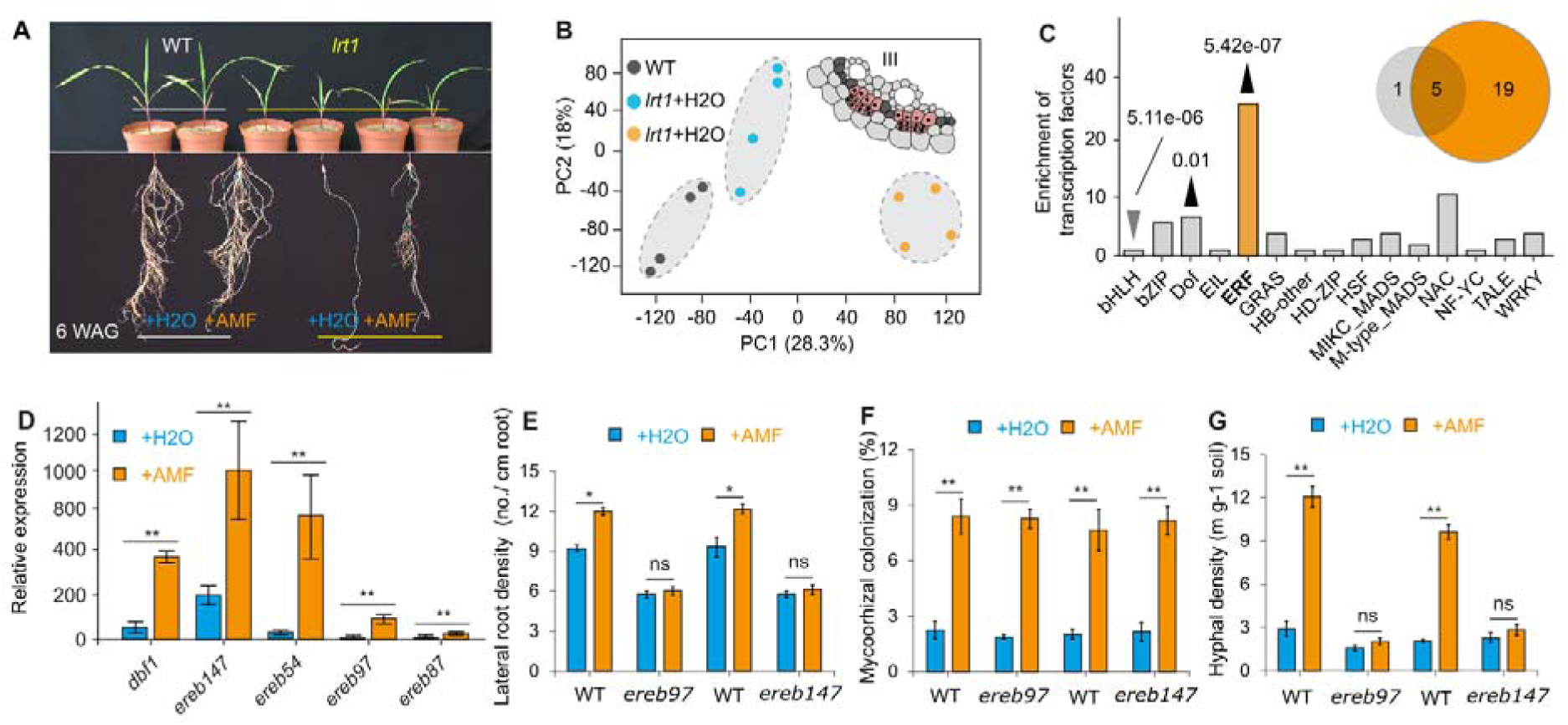
ERF transcription factors coordinate AMF-induced lateral root induction and rhizosphere hyphal expansion in maize. (**A**) AM inoculation recovered lateral root formation in the *lrt1* mutant. (**B**) AM inoculation largely reshaped the pericycle cell transcriptome. Here the stage III of the pericycle cells were captured and analysed by RNA-seq. (**C**) Enrichment analysis of transcription factors families. Statistical significance was controlled by chi-square test. An embedded Venn diagram indicates the overlapped transcription factors that AM fungi induced and the ones LRT1 silenced. (**D**) Relative expression of ERF genes in *lrt1* mutant induced by AM inoculation. Statistical significance was controlled by paired Student’s *t*-test. (**E**) AM mediated lateral root formation depends on function of ERF genes (erf mutant has no lateral root induction by AMF). (**F**) ERF mutation does not affect the AM fungal colonization, but hyphal branching in the rhizosphere (**G**).

Pairwise comparisons between *lrt1* roots with and without AM inoculation identified 1,855 differentially expressed genes (|FC| ≥ 2; FDR ≤ 0.05), with 1,004 genes upregulated upon AM colonization (Supplemental Fig. 4). Gene ontology (GO) enrichment highlighted “regulation of transcription, DNA-dependent” (GO:0006355) as the most significantly overrepresented biological process (FDR = 7.44e-07), pointing to transcriptional reprogramming as a central feature of AM-induced root remodeling. Notably, transcription factor family enrichment analysis revealed ERF (ethylene response factor) and Dof as the only significantly induced families (adjusted P = 5.42e-07 and 0.01, respectively; Fig. 2C). Among the AM-induced *ERF* genes, five overlapped with those downregulated in *lrt1* mutants, implicating them as key mediators of AM-triggered developmental rescue (Fig. 2D). Functional validation confirmed that AM fungi failed to induce lateral root formation or increase biomass in *erf* mutants, underscoring the necessity of *ERF* function for AM-mediated developmental plasticity (Fig. 2E). These results demonstrate that *ERF* gene activation is required for AM-induced lateral root development, particularly under low-phosphorus conditions.

To assess whether ERF function also affects AM fungal colonization, we quantified colonization rates in *erf* mutants and found no significant difference compared to wild-type roots (Fig. 2F), suggesting that while ERF activity promotes host root development, it is not essential for successful mycorrhizal establishment. Interestingly, mutation of *ERF* genes resulted in a significant reduction in hyphal length density in the rhizosphere (Fig. 2G), indicating that ERF-mediated root development may actively influence the extent of extraradical hyphal proliferation. This finding suggests a previously unrecognized role of host transcriptional regulation in shaping fungal foraging behavior and spatial colonization patterns in the soil environment. Together, these transcriptomic and genetic results establish *ERF* genes as critical integrators of AM fungal signaling and host developmental reprogramming, enabling lateral root induction and mutualistic symbiosis in maize.

### Cortex-localized flavonoid remodeling underpins AM-induced developmental plasticity

To further understand the metabolic signature between AM fungi and host root development, we employed integrative metabolomics (UPLC-MS/MS) and proteomics (iTRAQ) to profile tissue-specific responses in the stele and cortex of *lrt1* roots upon AM fungal colonization (Fig. 3A). Partial least squares discriminant analysis (PLS-DA) revealed a strong separation of metabolic profiles across tissues and treatment conditions, indicating tissue-specific and fungus-responsive metabolic reprogramming (Supplemental Fig. 6A). Quantitatively, 611 metabolites in the stele and 447 metabolites in the cortex were significantly altered (VIP ≥ 1; FC ≥ 1.2 or ≤ 0.8; *q* < 0.05) following AM inoculation in *lrt1* roots (Supplemental Fig. 6B). Interestingly, many of these metabolite levels were reduced, suggesting a potential scavenging or redirection of specific metabolite classes by the AM fungi (Supplemental Fig. 6C). Chemical similarity enrichment analysis (ChemRICH) performed on cortex-derived metabolites revealed significant enrichment of flavonoids and sesquiterpenes (Kolmogorov–Smirnov test < 0.05), both of which are known to play critical roles in plant–microbe communication (Fig. 3B). Multiple flavonoid subclasses—including flavonols (e.g., quercetin, kaempferol), flavones (e.g., luteolin), and their glycosylated or methylated derivatives—were identified and quantified (Fig. 3C). The enrichment of metabolites along the F3H–FLS–F3’5’H and 3/7-OMT branches of the pathway suggests that AM symbiosis alters flux through specific branches of the flavonoid biosynthetic network. Particularly, the consistent changes in quercetin-and luteolin-derived derivatives (e.g., 3’-O-methylquercetin, 3,7-O-dimethylquercetin, and luteoloside) imply that methylation and glycosylation steps are highly responsive to AM fungal cues. Pathway-level reconstruction using metabolic network analysis further emphasized the central role of flavonoid biosynthesis in cortex tissue responses to AM colonization (Supplemental Fig. 7; Supplemental Table 4). Correspondingly, proteomic analyses identified 163 differentially accumulated proteins in the cortex tissue of *lrt1* upon AM inoculation (Fig. 3D). Among these, 80 proteins were annotated via KEGG, with 19 significantly upregulated in the phenylpropanoid biosynthesis pathway (KEGG Map00940), a metabolic route upstream of flavonoid production. Notably, 14 of these genes showed cortex-specific expression, and 12 encoded class III peroxidases, enzymes involved in secondary metabolite processing and reactive oxygen species detoxification (Supplemental Table 5). The concurrent upregulation of peroxidases and flavonoid biosynthetic components suggests a tight coordination between redox regulation and specialized metabolite remodeling during AM-root symbiosis.

**Figure 3.**
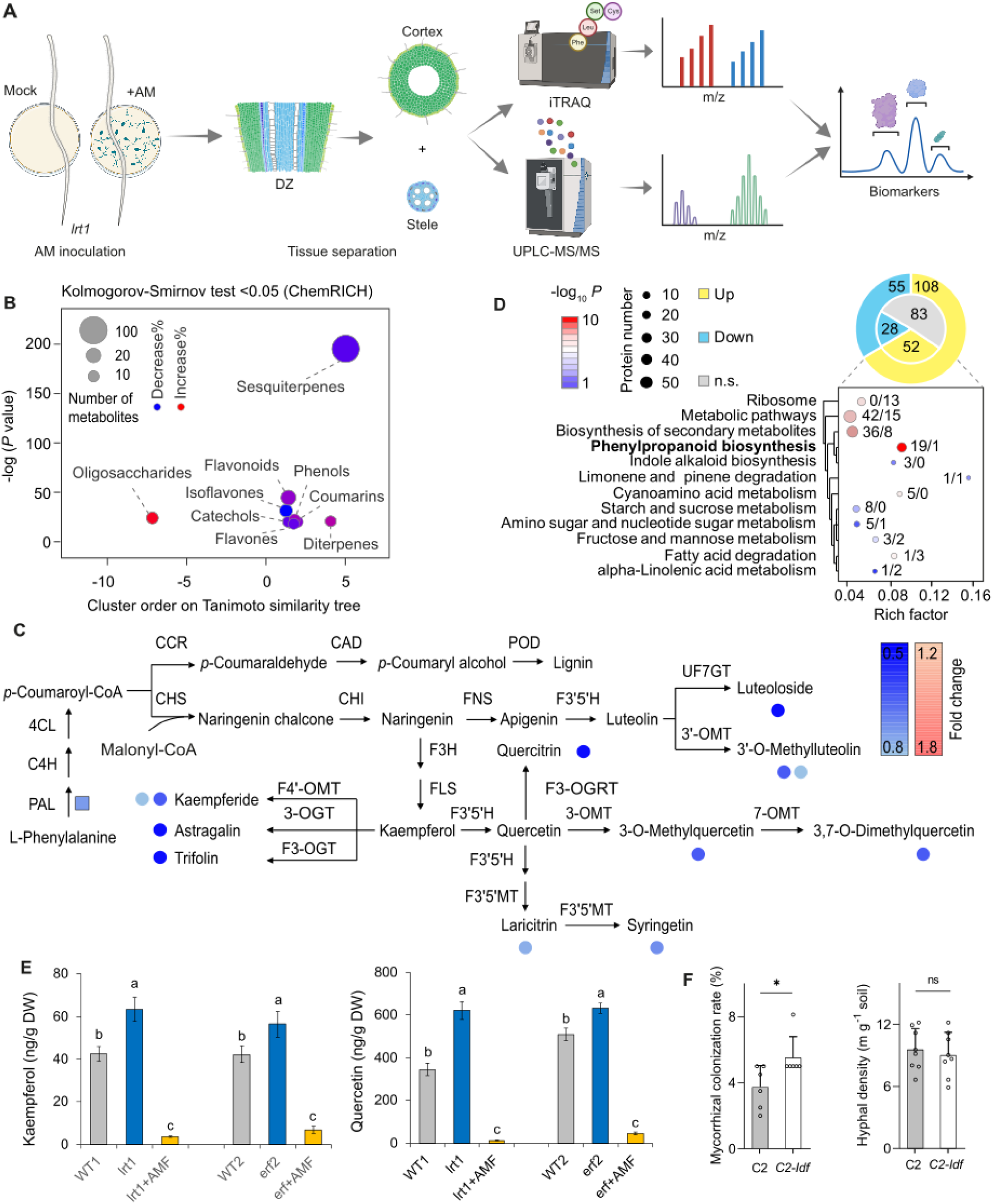
AM inoculation associated with host tissue-specific flavonoids homeostasis. (**A**) Schematic steps and workflow of root tissue-specific proteomic and metabolic analyses involved into AM symbiotic processes of *lrt1* mutant. (**B**) AM fungal inoculation significantly altered metabolite accumulation in the root cortex of *lrt1*, highlighting specific metabolic reprogramming in association with flavonoids. (**C**) AM symbiosis reshaped the flavonoid biosynthesis pathway, with differential regulation of ke enzymatic steps including CHS, CHI, FLS, F3H, and various O-methyltransferases and glycosyltransferases. 3-OGT: flavonol 3-O-glucosyltransferase; 3’OMT: flavone 3’-O-methyltransferase; 3-OMT: quercetin 3-O-methyltransferase; 4CL: 4-coumarate-CoA ligase; 7-OMT: 3-methylquercetin 7-O-methyltransferase; C4H: cinnamate 4-hydroxylase; CAD: cinnamyl alcohol dehydrogenase; CCR: cinnamoyl-CoA reductase; CHI: chalcone isomerase; CHS: chalcone synthase; F3’5’H: flavanoid 3’,5’-hydroxylase; F3’5’MT: flavonoid 3’,5’-methyltransferase; F3H: flavanone 3-hydroxylase; F3-OGRT: flavonol-3-O-glucoside L-rhamnosyltransferase; F3-OGT: kaempferol 3-O-galactosyltransferase; F4’-OMT: kaempferol 4’-O-methyltransferase; FLS: flavonol synthase; FNS: flavone synthase; PAL: phenylalanine ammonia-lyase; POD: peroxidase; UF7GT: flavone 7-O-beta-glucosyltransferase. (**D**) Proteomic analysis revealed significantly accumulated proteins and associated Phenylpropanoid biosynthesis pathway enrichments in response to AM colonization. (**E**) AM inoculation suppressed excessive flavonoid accumulation in both *lrt1* and *erf* mutants, indicating that the AM symbiosis modulates host secondary metabolism even in regulatory mutants. Statistical significance was assessed using one-way ANOVA, and different letters indicate significant differences among genotypes. (**F**) In contrast, the flavonoid-overproducing mutant C2 exhibited inhibited AM colonization despite high hyphal density in the rhizosphere, suggesting that excessive flavonoids may restrict fungal entry but not external fungal growth. Statistical significance was evaluated using a one-sided Student’s t-test. * indicates *P* value < 0.05, ns, not significant.

To investigate whether the ERF transcription factors modulate AM fungal–induced flavonoid metabolism, we quantified the major flavonoid compounds kaempferol and quercetin in both *lrt1* and *erf* mutants with or without AMF inoculation (Fig. 3E). Notably, in the absence of AM fungi, both *lrt1* and *erf* mutants accumulated significantly higher levels of kaempferol and quercetin compared to their respective wild-type controls, suggesting a feedback misregulation of flavonoid biosynthesis. Upon AM fungal inoculation, the concentrations of these flavonoids were dramatically reduced in both mutants, indicating that AM fungi may suppress flavonoid accumulation in roots lacking functional LRT1 or ERF, possibly due to impaired signal transduction or developmental competency in these backgrounds. To further dissect whether flavonoids themselves are required for AM fungal colonization, we assessed AMF symbiosis in a high-flavonoid-accumulating wild-type line (*C2*) and its flavonoid-deficient mutant (*C2-idf*). The *C2-idf* mutant showed a significant reduction in AM fungal colonization rate compared to *C2*, despite similar hyphal density in the rhizosphere soil (Fig. 3F). These results suggest that host-derived flavonoids facilitate successful fungal entry or establishment within roots, rather than merely affecting fungal proliferation in the surrounding soil. Together, these results demonstrate that ERF transcription factors and flavonoid metabolism form a critical regulatory hub at the AM fungi–maize root interface, coordinating both fungal colonization and lateral root development through transcriptional and metabolic reprogramming in the root cortex.

### Host-derived flavonoids gate selective *Massilia*–AMF interactions

To explore how host flavonoids modulate microbial associations in the rhizosphere, we analyzed the microbiome composition of the maize *lrt1* mutant, which lacks functional lateral root formation, and observed significant depletion of specific bacterial taxa such as *Massilia*, *Pseudomonas*, and *Sphingobium* compared to wild type (WT) roots, while bulk soil remained unaffected (Fig. 4A). This indicates a strong host-genotype–dependent effect on rhizosphere community assembly. We next investigated the spatial context of these interactions. To this end we employed a dual-compartment growth system to separate the root and hyphal compartments (Fig. 4B). This allowed us to test whether bacteria can associate directly with AM fungal hyphae independent of root proximity. Microscopy revealed that selected *Massilia* strains (R1485, I317, Ox6) physically attached to growing hyphae, while non-beneficial strains (I20, *E. coli*, or heat-killed bacteria) showed no colonization (Fig. 4C), demonstrating a degree of specificity in fungal–bacterial interactions.

**Figure 4.**
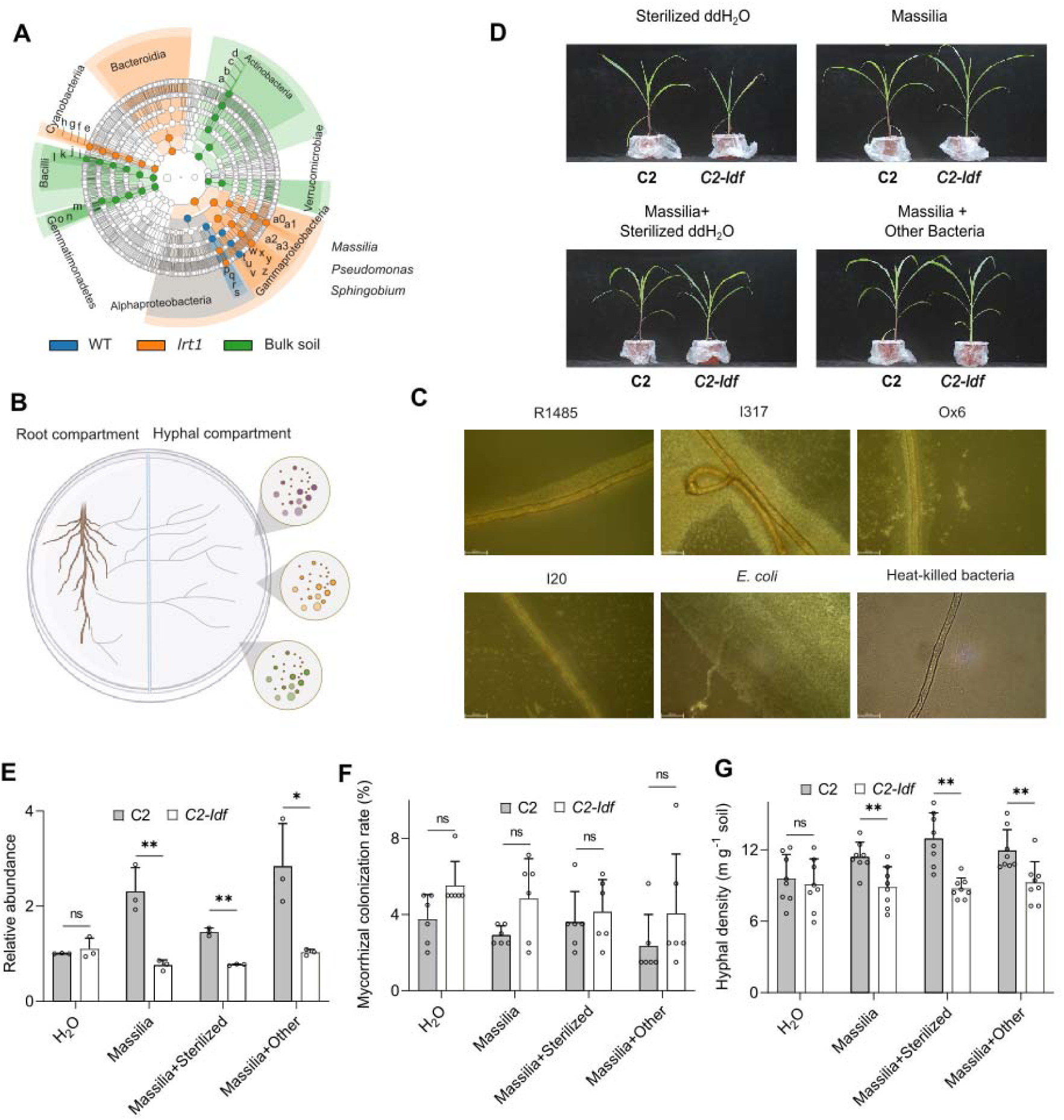
Flavonoids are required to maintain the AMF hyphae-associated enrichment of *Massilia* in the maize rhizosphere. **(A)** *Massilia* is selectively enriched in the rhizosphere of the *lrt1* mutant under low-phosphorus conditions, indicating a potential symbiotic association. **(B)** Schematic representation of an in vitro dual-compartment system used to study the interaction between AM fungal hyphae and *Massilia* colonization. **(C)** *Massilia* colonization along AM fungal hyphae visualized using a Leica Thunder Microscope. **(D)** Growth response of wild-type maize (C2) and its flavonoid-deficient mutant (*C2-Idf*) upon inoculation with either individual *Massilia* isolates or *Massilia*-based synthetic communities. **(E)** Host-derived flavonoids significantly enhance *Massilia* enrichment in the rhizosphere. **(F)** Mycorrhizal colonization rates remain unchanged between genotypes and treatments. **(G)** Flavonoids also promote increased AM hyphal length density in the rhizosphere.

We further tested whether host flavonoids are required for *Massilia*-mediated plant benefits. Using the flavonoid-rich maize genotype C2 and its flavonoid-deficient mutant *C2-idf*, we found that *Massilia* promoted plant growth and shoot biomass only in C2, but not in *C2-idf* mutant plants (Fig. 4D). This effect was not observed by applying heat-killed bacteria or co-inoculation with unrelated bacteria, underscoring the specificity of the C2–*Massilia* interaction. Quantification of *Massilia* abundance confirmed preferential recruitment in the rhizosphere of C2 plants relative to *C2-idf* (Fig. 4E). In line with this, *Massilia* significantly enhanced AM fungal colonization rates (Fig. 4F) and hyphal length density (Fig. 4G) in C2 but not in *C2-idf* plants. Neither heat-killed *Massilia* nor mixed bacterial communities reproduced these effects, further supporting the flavonoid-dependent nature of this tripartite interaction. These findings reveal that host-derived flavonoids act as selective gatekeepers mediating *Massilia* recruitment and AM fungal symbiosis, establishing a mechanistic link between root metabolites and beneficial microbiome assembly.

### Ethylene biosynthesis via ACS6/7 coordinates ERF activation and tripartite symbiosis

To integrate ethylene into the regulatory network, we analyzed *acs6* and *acs7* loss-of-function mutants and overexpression (OE,DPC) lines of the *ACS6* and *ACS7* genes, which encode key ACC synthases (Geisler-Lee et al., 2010). In wild-type maize, AMF inoculation significantly induced the expression of the ethylene-responsive transcription factors *ERF97* and *ERF147*. This induction was markedly reduced in *acs6* and *acs7* mutants, while *ACS6* and *ACS7* overexpression (OE) and promoter construct (PC) lines showed elevated expression of both ERFs, even in the absence of AMF (Supplemental Fig. 8A-B). These patterns suggest that ethylene biosynthesis via *ACS6/7* is required for AMF-induced ERF activation. Correspondingly, AMF-triggered increases in hyphal colonization and lateral root density were both impaired in *acs* mutants, but enhanced in OE and PC lines (Supplemental Fig. 8C-D). These findings support a model in which ethylene production mediated by *ACS6* and *ACS7* acts upstream of ERF-mediated transcriptional responses, facilitating both successful fungal colonization and the developmental reprogramming of root branching in response to AM symbiosis.

To dissect the role of ethylene biosynthesis in mediating such tripartite interactions, we employed *ACS7* overexpression (OE) lines and *acs7-1* loss-of-function mutants co-inoculated with *Massilia* and arbuscular mycorrhizal fungi (AMF). Phenotypic observations (Fig. 5A) revealed that *ACS7-OE1* plants showed enhanced shoot growth upon *Massilia* inoculation compared to wild-type (WT) and *acs7-1* lines. This benefit was attenuated when either sterilized *Massilia* or unrelated bacterial consortia were used. Quantification of AM fungal colonization (Fig. 5B) and hyphal density (Fig. 5C) showed no significant change across genotypes with *Massilia* alone, but significant increases were observed when *Massilia* was combined with other bacteria, highlighting a potential synergistic effect that depends on ACS7 activity. qPCR of *Massilia* abundance (Fig. 5D) further confirmed that *ACS7-OE* roots supported higher *Massilia* colonization than WT or *acs7-1*, suggesting ethylene biosynthesis modulates bacterial recruitment. Direct visualization of *Massilia* attachment to fungal hyphae in response to ACC supplementation (Fig. 5E) revealed enhanced hyphal colonization specifically in the hyphal compartment (HC) or when ACC was applied, indicating that host-derived ethylene precursors may influence fungal-associated bacterial enrichment. These effects were consistent across multiple *acs* alleles and transgenic lines (Fig. 5F), and positively correlated with increased lateral root density (Fig. 5G), shoot biomass (Supplemental Fig. 9A), and phosphorus accumulation in shoots (Supplemental Fig. 9B) under co-inoculated conditions. Collectively, these results establish that ACS6/7-dependent ethylene biosynthesis primes ERF activation, promotes beneficial *Massilia*–AMF consortia, and enhances nutrient acquisition at the maize root–soil interface.

**Figure 5.**
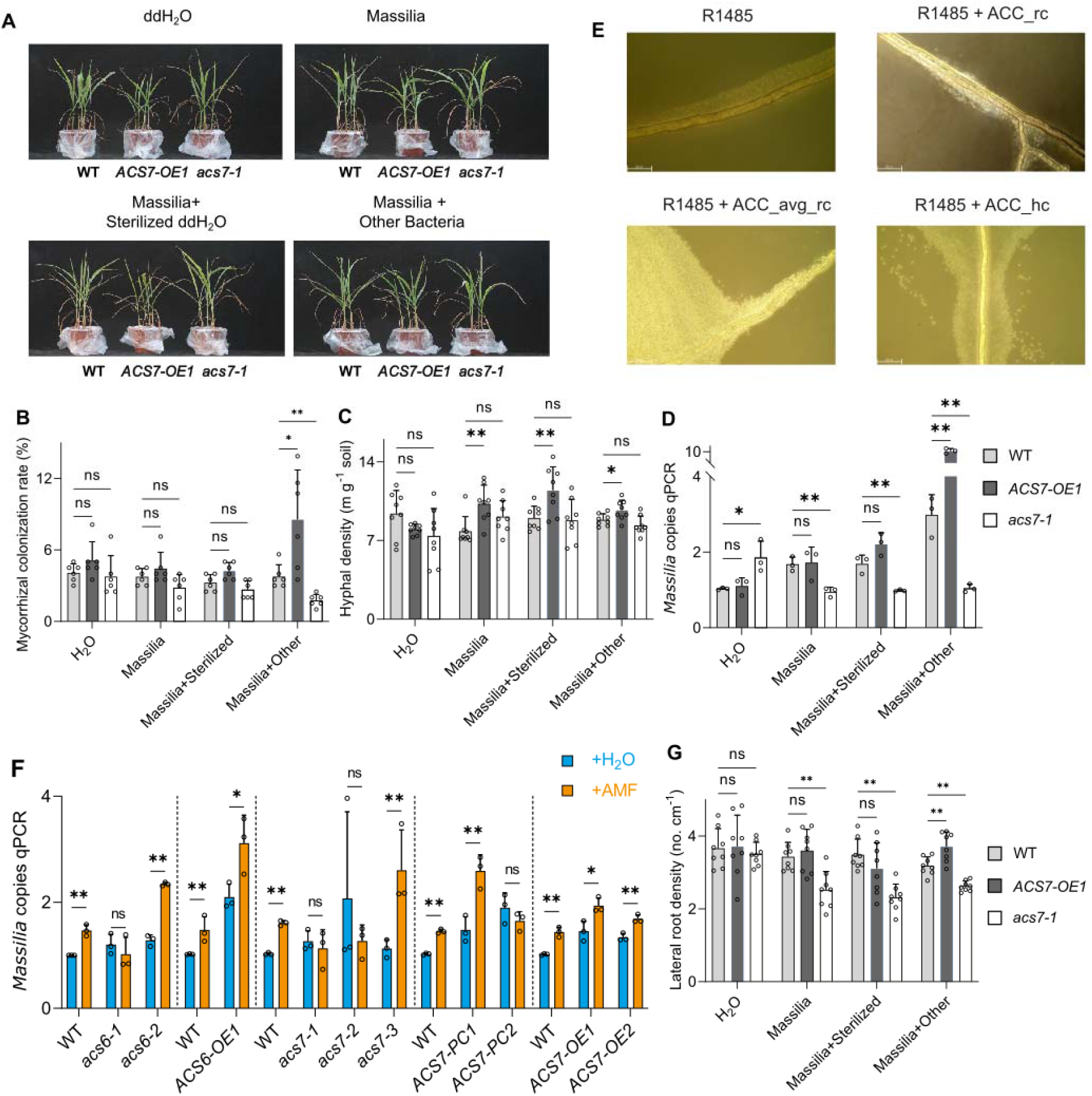
Ethylene mediates AM fungal cooperation with *Massilia* to enhance maize growth and phosphorus acquisition. **(A)** Growth responses of wild-type maize (B104), the ethylene biosynthesis mutant *acs7-1*, and overexpression line (ACS7-OE1) upon inoculation with either individual *Massilia* isolates or *Massilia*-based synthetic communities. **(B)** Mycorrhizal colonization rates are not significantly affected by genotype or *Massilia* treatment, indicating that ethylene does not alter AM fungal entry. **(C–D)** Ethylene biosynthesis is positively correlated with AM fungal hyphal length density (**C**) and *Massilia* enrichment in the rhizosphere (**D**) across treatments. **(E)** Supplementation of ACC (ethylene precursor) in the hyphal compartment enhances *Massilia* colonization along AM fungal hyphae, supporting ethylene-mediated hyphosphere recruitment. **(F)** AM fungal inoculation partially restores *Massilia* enrichment in ACS7 overexpression lines. **(G)** Ethylene biosynthesis is required for *Massilia*-induced lateral root development, as this response is impaired in *acs7-1* and enhanced in ACS7-OE1.

Taken together, across genetic, metabolic, and microbial analyses, our data revealed a coordinated regulatory network in which ethylenellldriven ERF activation modulates flavonoid homeostasis, enabling the assembly of AMF and *Massilia* at the root cortex. This tripartite integration of hormonal, metabolic, and microbial signaling defines a molecular framework for systematic engineering the plant–microbiome interface to improve plant productivity and resilience (Fig. 6).

**Figure 6.**
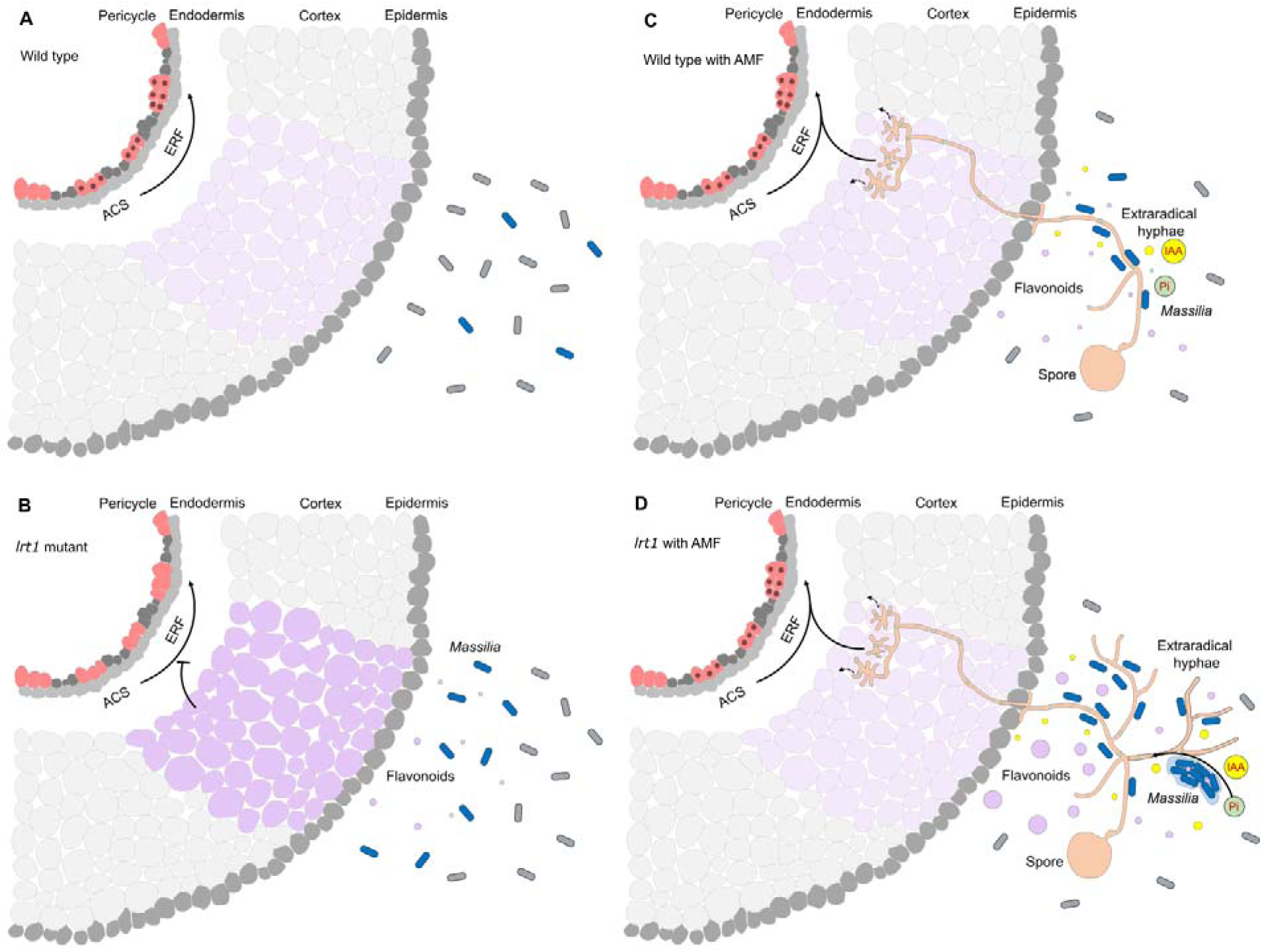
Conceptual model of ethylene pathway and flavonoid-mediated coordination of AM fungi, *Massilia*, and root development under low phosphate condition. This model illustrates how host-derived ethylene signaling integrates with flavonoid-mediated microbial recruitment to coordinate mutualistic interactions between arbuscular mycorrhizal (AM) fungi, *Massilia* bacteria, and root developmental pathways. Flavonoids act as key molecular signals connecting the plant, fungi, and bacteria, while also modulating ethylene biosynthesis. In turn, ethylene response factors (ERFs) regulate pericycle cell differentiation and lateral root formation, promoting enhanced nutrient uptake and adaptation to low-phosphorus environments. **(A)** In wild-type plants, the ZmACS–ZmERF signaling pathway regulates normal lateral root initiation, which occurs largely independently of rhizosphere bacterial microbiota. **(B)** In *lrt1* mutants, overaccumulation of flavonoids in the root cortex suppresses ERF gene expression, thereby inhibiting lateral root initiation. Although *Massilia* can still be partially recruited via flavonoid exudation, this is insufficient to restore normal root development. **(C)** In wild-type plants inoculated with AM fungi, extraradical hyphae associated with *Massilia*, promoting localized phosphorus solubilization and enhancing lateral root formation, providing an additional layer of symbiotic benefit. **(D)** In *lrt1* mutants colonized by AM fungi, cortical flavonoids are potentially redirected to the rhizosphere, where they enhance extraradical hyphal branching, promote *Massilia* enrichment, and facilitate biofilm formation on the hyphae. These interkingdom interactions stimulate auxin production and phosphoru solubilization, thereby partially restoring lateral root development despite disrupted host signaling.

## Discussion

### Interkingdom signaling at the root–soil interface: A coordinated role for root development, AM fungi, and beneficial bacteria

Our study uncovers a previously unknown coordination between root developmental programs and microbial partnerships in maize (Fig. 6). We demonstrate that the early stages of lateral root formation, particularly pericycle cell division, are intimately linked to the presence and activity of beneficial microorganisms, especially arbuscular mycorrhizal (AM) fungi and *Massilia* spp (Figs. 1, 2). In the lateral root-defective *lrt1* mutant, which is developmentally arrested before the emergence of lateral roots, AM fungi can partially restore lateral root formation, biomass accumulation, and root–soil interface functionality under phosphate-limiting conditions. This supports a model in which AM symbiosis not only facilitates nutrient uptake but also acts as a developmental modulator of host root architecture (Chiu et al., 2018, 2022; Zhang et al., 2024).

Consistent with previous findings (Sun et al., 2025; Wang et al., 2025), our dual-compartment and co-inoculation experiments show that *Massilia* spp, a beneficial rhizobacterium, selectively associates with AM fungal hyphae and enhances both hyphal proliferation and mycorrhizal colonization when host-derived flavonoids are present (Fig. 4). These findings suggest that specific root-associated bacteria can synergize with fungi to reinforce mutualistic functions, provided that the appropriate chemical and developmental context is established by the host. This aligns with recent reports that AMF do not act in isolation but rely on structured hyphosphere microbiomes—often enriched in bacterial taxa such as *Devosia*, *Streptomyces* and *Pseudomonas*—to optimize nutrient cycling and root colonization (Wang et al., 2022; Li et al., 2023; Jin et al., 2024; Zhang et al., 2024). Thus, the root–soil interface is shaped not merely by individual microbial taxa, but by a higher-order, interkingdom interaction between host developmental signals, fungi, and bacteria, forming a cooperative unit that supports sustainable plant growth.

### Flavonoids act as central chemical signals mediating tripartite symbiosis

Flavonoids have been recognized as pivotal signaling molecules in mediating arbuscular mycorrhizal (AM) symbiosis and also play a critical role in promoting nitrogen fixation by inducing nod gene expression in rhizobia, thereby orchestrating plant–microbe mutualisms across both fungal and bacterial domains (Wang et al., 2022; Bouwmeester et al., 2025). Several flavonoid compounds, such as daidzein, naringenin, and sakuranetin, have been shown to stimulate AM fungal spore germination (Scervino et al., 2005; Lidoy et al., 2023), enhance hyphal branching and extension (Buee et al., 2000; Scervino et al., 2006), and facilitate subsequent arbuscule formation within roots (Scervino et al., 2007). Building on this foundation, our multi-omics data revealed that AM fungi dynamically remodel flavonoid biosynthesis in maize roots, specifically targeting quercetin- and luteolin-derived compounds within the cortex (Fig. 3). These metabolite shifts suggest an active redirection of metabolic flux or fungal-mediated scavenging of host secondary metabolites. This aligns with emerging evidence that flavonoids not only serve as diffusible chemoattractants in the hyphosphere (He et al., 2025) but also function as selective gatekeepers that modulate microbial access to host tissues. In our system, flavonoid-deficient mutants (*C2-idf*) were unable to support AM fungal colonization or *Massilia* recruitment, despite normal hyphal density in bulk soil, underscoring the importance of flavonoid signaling for symbiotic specificity. Furthermore, we showed that the beneficial effects of *Massilia* on AM fungal function and plant performance are strictly dependent on host flavonoid biosynthesis. These findings highlight a refined role for flavonoids—not merely as growth-promoting signals, but as central chemical mediators of interkingdom communication, coordinating fungal entry, hyphal–bacterial interactions, and the downstream activation of root developmental programs under nutrient stress.

### ERF transcription factors integrate developmental and microbial signals in a unified regulatory hub

At the transcriptional level, ethylene-responsive factors (ERFs) represent a central regulatory node in coordinating root development (Cai et al., 2014) and fungal symbiosis cues (Das et al., 2025). Using laser capture microdissection, co-expression network analysis, and genetic mutants, we identified ERF genes (e.g., *ERF97*, *ERF147*) as critical drivers of pericycle division competence during lateral root initiation (Fig. 1). These ERFs were strongly induced during Stage III of lateral root formation and were significantly downregulated in *lrt1* mutants. Functional validation confirmed that ERF loss-of-function mutants display reduced lateral root density and fail to respond to AM-induced developmental cues. Importantly, we further demonstrate that ethylene biosynthesis genes (*ACS6*, *ACS7*) modulate ERF expression and are required for full AMF-induced root branching and *Massilia* colonization (Fig. 5). Overexpression of *ACS7* enhanced *Massilia* recruitment, hyphal density, and lateral root formation under AM conditions, while *acs7* mutants showed impaired responses. These results support a model in which ethylene biosynthesis acts upstream of ERF activation, which in turn orchestrates both developmental plasticity and microbial responsiveness. This finding is further supported by Gonin et al. (2023), who identified microbiota-induced ethylene response pathway as a central regulator of lateral root branching. Our findings establish that ERFs are not only master regulators of lateral root initiation but also key facilitators of multi-microbial symbiosis. By integrating signals from AM fungi, beneficial bacteria, and host hormone pathways, ERFs position themselves at the core of a regulatory network that controls root plasticity, microbial colonization, and nutrient acquisition. These findings are in line with growing evidence that host genetic components, such as transcription factors and metabolic enzymes, play a critical role in sculpting the root-associated microbiome (Stringlis et al., 2018; Yu et al., 2021; He et al., 2024).

## Conclusion

This study defines a mechanistic framework linking host developmental genes (LRT1, ERFs), hormonal pathways (ethylene via ACS6/7), and metabolic cues (flavonoids) to the recruitment and coordination of beneficial microbes (*Massilia*, AM fungi). Together, these results highlight the potential of engineering the root metabolite–microbiome–development axis to optimize plant–microbe interactions for improved crop performance under nutrient stress.

## Resource availability

### Lead contact

Requests for further information and resources should be directed to Peng Yu.

## Materials availability

Unique genetic resources and mutants generated in this study are available from the lead contract upon reasonable request.

## Data and code availability

- Raw plant LCM cell RNA-seq data reported in this paper were deposited in the Sequence Read Archive (http://www.ncbi.nlm.nih.gov/sra) under accession no. PRJNA1366082.
- This paper does not report any original code.
- Any additional information required to reanalyze the data reported in this paper is available from the lead contract upon request.

## Acknowledgements

We would like to thank Joachim Hamacher from the Institute of Crop Science and Resource Conservation (INRES), University of Bonn, for support in transmission electronic microscopy. We gratefully acknowledge the Center for Crop Functional Genomics and Molecular Breeding at China Agricultural University (CAU) for providing the ZmACS6 and ZmACS7 mutant and overexpression maize lines used in this study. This work is supported by the Deutsche Forschungsgemeinschaft (DFG) Emmy Noether Programme 444755415 to P.Y., research grants 514003603 and 552843484 to P.Y., the DFG Priority Program (SPP2089) ‘Rhizosphere Spatiotemporal Organisation - A Key to Rhizosphere Functions’ grant 403671039 to F.H. and P.Y., research grant HO 2249/12-2 to F.H. and the research group “FOR 5235: Cereal Stem Cell Systems (CSCS): Establishment, Maintenance and Termination” grant HO 2249/23-2 to F.H. This work was supported by the Biological Breeding-National Science and Technology Major Project (2024ZD04077) to X.L. Maize transformation was funded by Biological Breeding-National Science and Technology Major Project (2024ZD04077).

## Author contributions

Conceptualization, P.Y. and F.H.; coordination and supervision, P.Y.; methodology, M.B., Y.Z., B.Y., L.G., X.H., Ma. B., N. von W., C.G. and P.Y.; investigation, M.B., Y.Z., B.Y., Y.A.T.M., S.W., Z.Z., L.Z., L.Y., and P.Y.; bioinformatics, T.T., F.H. and P.Y.; genetic resources, R.S. and X.L.; writing, C.L., X.L., F.H., and P.Y.; funding acquisition, X.L., F.H., and P.Y.

## Supplemental information

Document S1. Figures S1–S8 and Table S1-S5

## Materials and Methods

### Plant Material and Germplasm Resources

The monogenic recessive maize root mutant *lrt1* (*lateralrootless 1*, Hochholdinger and Feix, 1998) and its corresponding wild type B73 were used. Moreover, homozygous naturally silenced maize *C2-Idf* (*Colorless2-inhibitor diffuse*) mutants and corresponding C2 wild type maize seeds were analysed (Della Vedova et al. 2005). In addition, we used recombinant inbred line (RIL) population derived from four generations of intermating of the inbred lines B73 and Mo17 (IBM) followed by seven rounds of selfing (Lee et al. 2002). Of the 163 RILs, 12 lines displayed a significantly distinct lateral root density. Among those the inbred lines 2033, 2043, 2156, 2215, 2242, 2274 displayed a low lateral root density) and the inbred lines 2088, 2104, 2161, 2162, 2168, 2236 displayed a high lateral root density. To generate the construct of CRISPR/Cas9-mediated knockout lines of *ZmACS6* and *ZmACS7* and perturbation of the C-terminal domain (PC) of *ZmACS7* editing, the specific 19-bp sgRNA (GTCGGAGCAACGTGGGTAC, TAAATGGCCGGTAGCAGCG, GAGATGGCGAGATGCGAAG) was cloned into the pBUE411 vector, respectively. To construct the overexpression lines of *ZmACS6* and *ZmACS7*, their full-length coding sequences were amplified from the cDNA of B73 and cloned into the pBCXUN vector to generate the *Ubi:ZmACS6* and *Ubi:ZmACS7* constructs. The knockout, PC, and overexpression lines of *ZmACS7* were used as previously described (Li et al., 2020). All seeds of the *ZmACS6* and *ZmACS7* transgenic lines were kindly provided by the Center for Crop Functional Genomics and Molecular Breeding of China Agricultural University. All primers were listed in Table S6. To the end, CRISPR-Cas9 knock out lines were generated for the maize genes *Zmereb97* and *Zmereb147* using the previously described simplex strategy (Qi et al. 2016). As a result the 47 bp sequence 5′ CCG AGG TAC CGC GGC GTC CGG AAG CGC CCC TGG GGA CGC TAC GCG GC 3′) sequence was deleted in the protein-coding region of *Zmereb97* and a 1 bp adenine was inserted in *Zmereb147*. The *Agrobacterium*-mediated maize transformation procedure was performed as described in the simplex strategy (Qi et al. 2016).

### Experimental Set-up and Growth Conditions

For the phytochamber experiments, surface-sterilized seeds of the mutants *lrt1* and *C2-idf* and their corresponding wild type controls described above have been pre-germinated for three days in a paper roll system with deionized water in a 16-h-light, 26 °C, and 8-h-dark, 18 °C cycle. Germinated mutants and wild type seedlings with 2-3 cm length primary roots were transferred to new paper rolls and randomly distributed in a 5L beaker supplied with air and grown for another seven days. The same germination and cultivation procedures were applied to the 163 maize RILs used for phenotyping lateral root density.

For soil cultivation and inoculation with *R. irregularis*, 3-day pre-germinated mutants and wild type seedlings were transferred into an artificial soil containing sterilized coarse sand, fine quartz and loam (8:1:1 by weight) in a 2 L pot. For inoculation with *R. irregularis* (quality A spores from Agronutrition, Carbonne, France), the spore stock (2,000 spores per ml) were placed in a 50 ml Falcon tube and centrifuged at 3,000 x *g* for 10 min to spin down the spores. The supernatant (spore germination inhibition buffer) was removed and the spores were washed three times with autoclaved AM water (50% tap water and 50 % deionized water) at 3,000 x *g* for 10 min. The final inoculants with 1,000 spores per plant were applied to the soil upon planting the 3-day-old seedlings. The plants were supplemented with modified Hoagland solution (Yu et al. 2015) with reduced phosphate (0.1 mM KH_2_PO_4_) and were grown 4-6 weeks in a well-controlled phytochamber with a 16-h-light, 26 °C, and 8-h-dark, 18 °C cycle.

For collecting the germinated AM fungal spores exudates, the stock spores in sterile AM water were placed in petri dishes to generate a large surface for air exchange and germinated at 30 °C for 14 days. During that time, the solution medium was corrected to the original volume due to evaporation. The germinated spores were checked under the stereomicroscope (Leica M165 FC) to confirm hyphae formation. The germinated spores were then centrifuged as described above and the supernatant of the germinated spore exudates were used to treat the recipient plants.

### Soil Pot Experiments for Maize Ethylene Mutants

For soil culture experiments, the moderately acid soil (pH=6.36) which contained 4.24 mg olsen phosphorus (P) kg^−1^, 0.29 g total nitrogen (N) kg^−1^, and 27.02 mg exchangeable potassium (K) kg^−1^ was collected from Shangzhuang, Haidian, Beijing, China. Maize seeds of inbred lines B73-329 (WT), *ZmACS6* knockout (KO) and overexpression (OE) lines, *ZmACS7* (KO) knockout, overexpression (OE), and gain of function lines (PC) were germinated and grown in a pot with 1.8 kg of the low phosphorus soil, premixed with 200 mg kg^−1^ (NH_4_)_2_SO_4_, 40 mg kg^−1^ KH_2_PO_4_, 200 mg kg^−1^ K_2_SO_4_, 50 mg kg^−1^ MgSO_4_, 5 mg kg^−1^ ZnSO_4_, 50 mg kg^−1^ MnSO_4_, 5 mg kg^−1^ CuSO_4_. 2 mL of the *R. irregularis* spore solution (42 spores mL^−1^) was added to the soil for AM fungus inoculation treatment and the same volume sterilized water for the no-inoculation treatment. The seedlings were harvested after 8 weeks of soil culture. Crown root samples and the rhizosphere soil were collected and stored at −20°C and −80°C freezers for further analysis.

To further determine whether *Massilia* can promote maize growth under low phosphate conditions with AMF inoculation. Inbred lines B73-329 (WT), *zmacs7-1*, *ZmACS7-OE1*, *C2*, and *C2-idf* were inoculated with isolated strains including seven *Massilia*, thirteen *Telluria*, three *Collinonas*, two *Duganella*, and one *Pseudoduganella*. All strains were cultured overnight at 28°C in R2A liquid medium and then centrifuged for 10 min at 1,500 g. The pellets were resuspended in sterilized water and diluted to an optical density OD600 = 0.02, respectively. Before the inoculation experiment, the soil was sterilized with ^60^Co γ-radiation to kill the soil-borne microbes with a dose of 20 kGy at Beijing Atomic High-tech Jinhui Radiation Technology Application Co. Ltd. Four treatments were set up, including sterile water, inoculation with the *Massilla* strain, inoculation with the *Massilla* strain and sterile water, and inoculation with the *Massilla* and nineteen other mixed strains treatments. We harvested the plants and soil samples at 8 weeks after soil culture for further analysis.The shoot samples were harvested for dry weight and P content analysis.

### Dry Weight and P Content Measurement

The shoot samples were oven-dried at 105°C for 30 min to stop metabolic activity and then dried at 65°C until constant weight. After weighing the dry weight by using a balance, the samples were ground into a powder and digested using H_2_SO_4_ and H_2_O_2_. Then digests were analyzed for P content by the spectrophotometric colorimetric method (Soon and Kalra, 1995).

### Morphological, Histological and Anatomical Analyses of Lateral Root Initiation

To determine lateral root density, primary roots of the mutants and the wild type plants together with the RILs were dissected and the total emerged lateral root number was hand-counted. Lateral root density was calculated as the number of emerged lateral roots divided by the length of the region where lateral roots emerged.

Developmental stages of lateral root primordia were quantified from 4 cm long root segments from the root tip of *lrt1* mutant seedlings and its wild type. In brief, 4 cm long primary roots were dissected into 5 mm segments. These segments were then fixed and 10 µm longitudinal paraffin sections were prepared by a rotary microtome (Leica) as described previously (Lim et al. 2000). Deparaffinized specimens were gradually dehydrated then subjected to a acid-hydrolysis reaction with freshly prepared 1N HCl at room temperature for 30 min. The hydrolyzed specimens were immediately dried with filter paper and stained by Schiff’s solution (Sigma) for 1 hour in the dark. The stained specimens were directly transferred to a freshly prepared bleach solution (1N HCl, 10% K_2_S_2_O_5_ and water with 1:1:18 ratio by volume) and were then counterstained with Toluidine Blue (0.01% in water for 10-30 sec at room temperature). The stained material wasthen dehydrated and embedded in Entellan (Merck) and analyzed under an AxioCam MRc Microscope (Carl Zeiss Microimaging) and documented with Axio-Imager software (Carl Zeiss Microimaging). The frequency of lateral root initiation was illustrated as the lateral root initiation index (Dubrovsky et al. 2009) and lateral root primordia were further classified according to their developmental stages (Malamy and Benfey 1997).

### Isolation of Pericycle Cell Types by Laser Capture Microdissection

To understand the early events during lateral root initiation, pericycle cells adjacent to phloem and xylem poles were isolated from cryosections as previously described (Yu et al. 2015). In brief, root segments at a distance of 5-10 mm (Segment 1), 10-15 mm (Segment 2) and 15-20 mm (Segment 3) from the primary root tip of *lrt1* and wild type seedlings were dissected and fixed immediately in a freshly prepared farmer’s fixative. After a series of vacuum infiltration/swirl and protecting/swirl steps, the root segments were embedded in Tissue Freezing Medium and stored at −80°C until cryosectioning. The embedded molds with root segments were cryosectioned in 20 µm sections using a Cryotome (Leica) at −28°C. The trimmed cryosections were then mounted with PEN Membrane Slides (Carl Zeiss Microscopy). Prior to laser capture microdissection (LCM), the tissue freezing medium was removed by sequential steps of dehydration with ethanol and xylene solutions. LCM was performed on the air-dried sections with a PALM MicroBeam Platform (Carl Zeiss Microimaging). Approximately ≥20 phloem-pole or xylem-pole pericycle cells were isolated from each section and in total ca. 5,000 cells were pooled for each cell-type from respective 5 mm root segment for downstream analysis. For the soil AM inoculation experiments, the same LCM work pipeline was applied for crown roots of 4-week old *lrt1* plants with or without AM inoculation. In particular, only phloem-pole pericycle cells were isolated from each section and in total around 15,000 cells were pooled from respective 15 mm root segments for each mycorrhiza treatment.

### Physical Separation of Root Stele Tissues from IBM RILs

The stele tissue contains the pericycle and is therefore enriched for cells involved in lateral root initiation. To identify genes associated with lateral root formation, the stele tissues were manually separated from the cortical parenchyma for the six RILs with high lateral root density and the six RILs with low lateral root density (Reference). In brief, the first 5 mm of the root tip was removed from nine-day-old seedlings grown in the paper roll system. Then the stele was separated manually from the cortical parenchyma in the root region until the first lateral root emerged. Dissected stele tissues of 6-8 cm length were pooled from the primary roots of ≥10individual plants for each RIL and stored at at −80 °C until RNA extraction.

### Isolation of RNA, cDNA Library Construction and Next Generation Sequencing

RNA extraction from the LCM captured cells and the separated root steles were performed as previously described (Yu et al. 2015, 2016). High quality RNA with RIN (RNA integrity number) values ≥6.5 were used for downstream amplification and cDNA synthesis as previously described (Yu et al. 2016). cDNA library construction, quantification and qualification of the sample libraries for Illumina sequencing were performed as previously described (Yu et al. 2016). The raw sequencing reads were obtained by Illumina sequencing. Subsequently, adapter sequences were trimmed and the quality score was checked using the FastQC program. High quality sequencing reads were processed and subsequently mapped to the maize B73 reference genome sequence (RefGen_v3) with CLC Genomics Workbench software. Only reads uniquely mapping to the reference genome were further analyzed. Prior to statistical analyses, data were normalized by calculating FPKM (fragments per kilobase of exon per million fragments mapped) values by dividing the total number of read pairs per gene by the product of mapped reads (millions) and exon length (in kb). Only genes that were represented by a minimum of five mapped reads in all four replicates of at least one cell type per segment per genotype were declared as expressed and considered for downstream analyses. The raw sequencing reads were normalized by sequencing depth and were log_2_-transformed to meet the assumptions of a linear model.

### Bioinformatics, Statistical Analyses of Sequencing Reads and Functional Characterization

The homogenous distribution of replicated samples and thus the quality of the sequencing output of the RNA-seq samples were determined by a principle component analysis (PCA) and hierarchical clustering by CLC genomics workbench. The cell transcriptome data for two pericycle cell types across the three segments between *lrt1* and its wild type were subjected to weighted gene co-expression network analyses (WGCNA) to determine the gene co-expression modules across the 48 genotype-segment-cell type-specific samples. Normalization, transformation and module detection was analysed as previously described (Yu et al. 2020). Annotation of co-expressed genes within modules was performed by AgriGO. Moreover, Fisher’s exact test (Du et al., 2010) was applied to determine significant enrichment of transcription factor families associated with the lateral root initiation. Furthermore, the CentriMo utility in MEME suite (Bailey et al. 2009) was used to perform transcription factor binding site enrichment analysis for WGCNA co-expressed module genes. We focused on the promoter regions as 1 kb upstream and 500 bp downstream of transcription start sites of co-expressed module genes after obtained the genomic sequences using a customized script. Within each module, 10 motifs with E-value less than 10^−6^ were recorded by MEME. Using the TOMTOM motif comparison tool, the resulting motifs were aligned with motifs in the JASPAR CORE Plants database (Mathelier et al. 2014) to identify significantly similar known *cis*-motifs (*q*-value < 0.05).

To identify the expression profile correlation and clustering among RILs with distinct lateral root densities, K-means clustering was performed with the K-means support module embedded in the MEV program with the figure of merit assessing the quality of the clustering algorithm (Yeung et al. 2001). Annotation of clustered genes was analyzed by AgriGO and association networks of the identified genes significantly enriched in GO terms were constructed with the aid of the online analysis tool STRING v10 (Szklarczyk et al. 2015).

To explore gene expression patterns after AM fungal inoculation, differential gene expression with or without AM fungal inoculation was performed by pairwise comparison for *lrt1* using the BIOCONDUCTOR package LIMMA (Smyth 2004) in R (v.3.1.1). Subsequently, we determined the number of differentially expressed genes for each comparison by controlling the *p*-values of the pairwise *t*-tests by adjusting FDR at 0.05 for multiple testing and a fold change |Fc| >2 (Benjamini and Hochberg 1995). Then, we functionally classified differential gene expression patterns according to Gene Ontology (GO) terms. The GO annotation system is based on three structured vocabularies that describe gene products in terms of their associated biological processes, cellular components and molecular functions in a species-independent manner. Subsequently, we performed a singular enrichment analysis to discover significantly overrepresented functional categories.

### Integrated Proteome and Metabolome Analyses of Root Stele and Cortex with AM Inoculation

An integrated analysis of multiple types of omics data may improve the understanding of the molecular mechanism of root symbiosis with AM fungi. We first applied ultra-performance liquid chromatography-tandem mass spectrometry (UPLC-MS/MS)-based metabolomics to quantify the untargeted metabolites in AM-inoculated *lrt1* plants. Metabolites were extracted from an amount of 25 mg frozen tissue samples with 800 µL cold 50% methanol. These extracts were then fully lysed in a Tissue Lyser (40Hz, 5min). The lysed samples were precipitated at −80 °C for 2 h, then the extracts were centrifuged at 25,000×*g* for 15 min at 4 °C and 550 µL supernatant was transferred to a new vial. Prior to the UPLC/MS run, the quality control samples were used to calibrate the UPLC column. The quality control samples were also injected regularly throughout the run (after every 10 samples) to monitor the stability of the analytical platform. A high-resolution tandem mass spectrometer SYNAPT G2 XS QTOF (Waters, USA) was used for metabolite detection with ten biological replicates. Metabolic features alignment, peak extraction, normalization and compound identification were performed by Progenesis QI (version 2.2). Moreover, quality control - based robust LOESS (locally estimated scatterplot smoothing) signal correction (QC-RLSC) was applied to efficiently correct the instrumental variation during metabolome analysis (Dunn et al. 2011). By combining univariate and multivariate statistical analysis, significantly changed features (FDR <0.05, VIP ≥1, FC ≥1.2 or ≤0.8) were acquired and those significant features were further annotated with the databases PubChem and KEGG. In addition, partial least squares - discriminant analyses (PLS-DA) were performed to maximize the covariance of metabolic features of tissues upon inoculation with AM fungi.

To investigate the effects of AM inoculation on the metabolism of stele and cortex, isobaric tags for relative and absolute quantitation (iTRAQ)-based proteome analysis was employed in parallel to investigate proteomic changes of AM inoculated roots. Approximate 2 g frozen root tissues with 10% polyvinylpolypyrrolidone were grounded into powder in liquid nitrogen and then sonicated on ice for 5 min in lysis buffer (8 M urea and 40 mM Tris-HCl containing 1 mM phenylmethylsulfonyl fluoride, 2 mM EDTA and 10 mM DTT, pH 8.5). Following centrifugation (25,000 *g*, 4 °C, 20 min), the supernatant was treated by adding 5 volume of 10% trichloroacetic acid/acetone with 10 mM DTT to precipitate proteins at −20 °C for 2 hours. The supernatant was discarded and the resulting pellet was washed twice with cold acetone on ice. Prior to protein digestion, the concentrations of extracted proteins were quantified by a Bradford assay using Coomassie Brilliant Blue G-250. Protein integrity was further determined by SDS-Polyacrylamide Gel Electrophoretogram. Protein digestion was performed with Trypsin Gold (Promega, Madison, WI, USA) at 37 °C overnight, followed by desalting of the peptides with a Strata X C18 column (Phenomenex) and vacuum-drying. Afterwards, the peptides were labelled with the iTRAQ Reagent 8-plex Kit according to the manufacturer’s protocol (Applied Biosystems). The iTRAQ-labelled peptides were subjected to strong cation exchange fractionation on a Shimadzu LC-20AB HPLC Pump system coupled with a high pH RP column. The peptides were reconstituted with buffer A (5% ACN, 95% H_2_O, pH 9.8) to 2 ml and loaded onto a column containing 5 μm particles (Phenomenex). The peptides were separated at a flow rate of 1 mL/min with a gradient of 5% buffer B (5% H_2_O, 95% ACN, pH 9.8) for 10 min, 5-35% buffer B for 40 min, 35-95% buffer B for 1 min. The system was then maintained in 95% buffer B for 3 min and decreased to 5% within 1 min before equilibrating with 5% buffer B for 10 min. Elution was monitored by measuring absorbance at 214 nm, and fractions were collected every 1 min. The eluted peptides were pooled as 20 fractions and each fraction was concentrated via vacuum centrifugation. The iTRAQ-labelled samples were analyzed using a LC-20AD nano-HPLC instrument (Shimadzu, Kyoto, Japan). Then, the peptides were eluted from trap column and separated by an analytical C18 column (inner diameter 75 μm) packed in-house. Data acquisition was performed with a TripleTOF 5600 System (SCIEX, Framingham, MA, USA) equipped with a Nanospray III source (SCIEX, Framingham, MA, USA), a pulled quartz tip as the emitter (New Objectives, Woburn, MA) and controlled with software Analyst 1.6 (AB SCIEX, Concord, ON). The raw MS/MS data was converted into MGF format by ProteoWizard tool msConvert, and the exported MGF files were searched using Mascot (version 2.3.02) against maize UniProt protein database (https://www.uniprot.org/proteomes/UP000007305). IQuant (Wen et al. 2014) was applied for quantitatively analyzing the labelled peptides with isobaric tags. All differentially accumulated proteins with a false discovery rate (FDR) >1% were included in downstream analyses including Gene Ontology (GO) enrichment analyses, KEGG pathway analyses and cluster analyses.

### Quantification of AM Colonization and Microscopic Imaging, and Hyphal Density in Soil

Representative primary and crown roots were collected from *lrt1* and its wild type plants 6 weeks post-inoculation with *G. intraradices*, along with crown roots from the mutants of *ZmACS6* and *ZmACS7* and the corresponding B73-329 wild type lines after 8 weeks of inoculation with *R. irregularis.* Roots were stained with trypan blue and mycorrhizal colonization was quantified with a modified grid-line intersect procedure (Paszkowski et al. 2006). In brief, the total root length colonization was determined by the total amount of fungus scored microscopically at 100 random points per root sample. We also performed the trypan blue staining to determine the hyphal length density in rhizosphere soil according to the previous method (Brundrett et al., 1994).

### RNA Extraction and Quantitative real-time PCR

RNA extraction, cDNA synthesis and real time RT-PCR were performed as previously described (Yu et al. 2015). RNA was isolated from 100 mg of liquid nitrogen-ground root samples with Trizol reagent according to the manufacturer’s instruction (Invitrogen). The quality and quantity of isolated RNA were determined using a NanoDriop Spectrophotometer (Thermo Fisher Scientific, Waltham, MA, United States). 1 g of total RNA was purified and reverse transcribed into cDNA using the PrimeScript™ RT reagent Kit (Takara, Kyoto, Japan). TB Green® Premix Ex Taq™ (Takara, Kyoto, Japan) was used to detect the relative gene expression with *ZmTublin* as the internal control by RT-qPCR in a CFX96 Touch Real-Time PCR Detection System (Bio-Rad). All primers were listed in Table S6.

### DNA Extraction and RT-qPCR Analysis of *Massilia* Relative Abundance

The DNA extracted from 0.5 g rhizosphere soil was performed using FastDNA™ Spin Kit for Soil DNA Extraction followed by instructions (MP Biomedicals, Irvine, CA, United States). Then the relative abundance of *Massilia* was quantified by RT-qPCR with the universal bacteria 16S rRNA gene and *Massilia* specific 16S rRNA gene. The primers were 1048F (5’-GTGSTGCAYGGYTGTCGTCA-3’)/1194R (5’-ACGTCRTCCMCACCTTCCTC-3’) and F (5’-GGAGCGGCCGATRTCTGATTAG-3’)/R (5’-ATTCTACCCCCCTCTGCCARAY-3’) for the universal bacteria 16S rRNA gene and *Massilia* specific 16S rRNA gene, respectively.

### TEM Microscopy

Primary root of *lrt1* with or without inoculation by *G. intraradices* were fixed in Karnoskýs fixative solution (Karnovsky 1965) at room temperature for 2 h. Washing steps using cacodylate buffer were repeated for 10 times and each time for 10 min, followed by staining with 1% osmium tetraoxide solution for 1 h then washed again with the same buffer for 10 times. Dehydration steps were performed with ethanol series from 15%, 20%, 50%, 70% and 90% for 10 min, two times of pure ethanol for 30 min and two times 99.5% propylene oxoide for 10 min. Afterwards, the root samples were embedded with gradual concentrated low viscosity Spurr resin (Sigma-Aldrich) with different percentage of propylene oxoide with 1:3, 1:1, 3:1 and pure Spurr resin for 12 hours each. Then, the samples were polymerized in pure Spurr resin in rubber embedding trays at 70°C for 16h. For sectioning, 70-80 nm thickness of resin sections were generated with Reichert-Jung Ultramicrotome Ultracut E (Leica Microsystems). The sections were stained with fresh 2% in aequous solution of uranyl acetate for 2 min, washed with bidistilled water and subsequently stained with Reynolds lead citrate under CO_2_ reduction with NaOH pellets for 20 sec, washed again and dried on forceps before examination. Transmission electron microscopy measurements were carried out on a Zeiss electron microscope 109T and the images were obtained with a slow-scan CCD camera from TRS (Tröndle, Moorenweis, Germany).

