## Supplemental figures and tables for "Ethylene signal-driven plant-multitrophic synergy boosts crop performance" (from another version of this preprint: version 1, 2025-11-29)

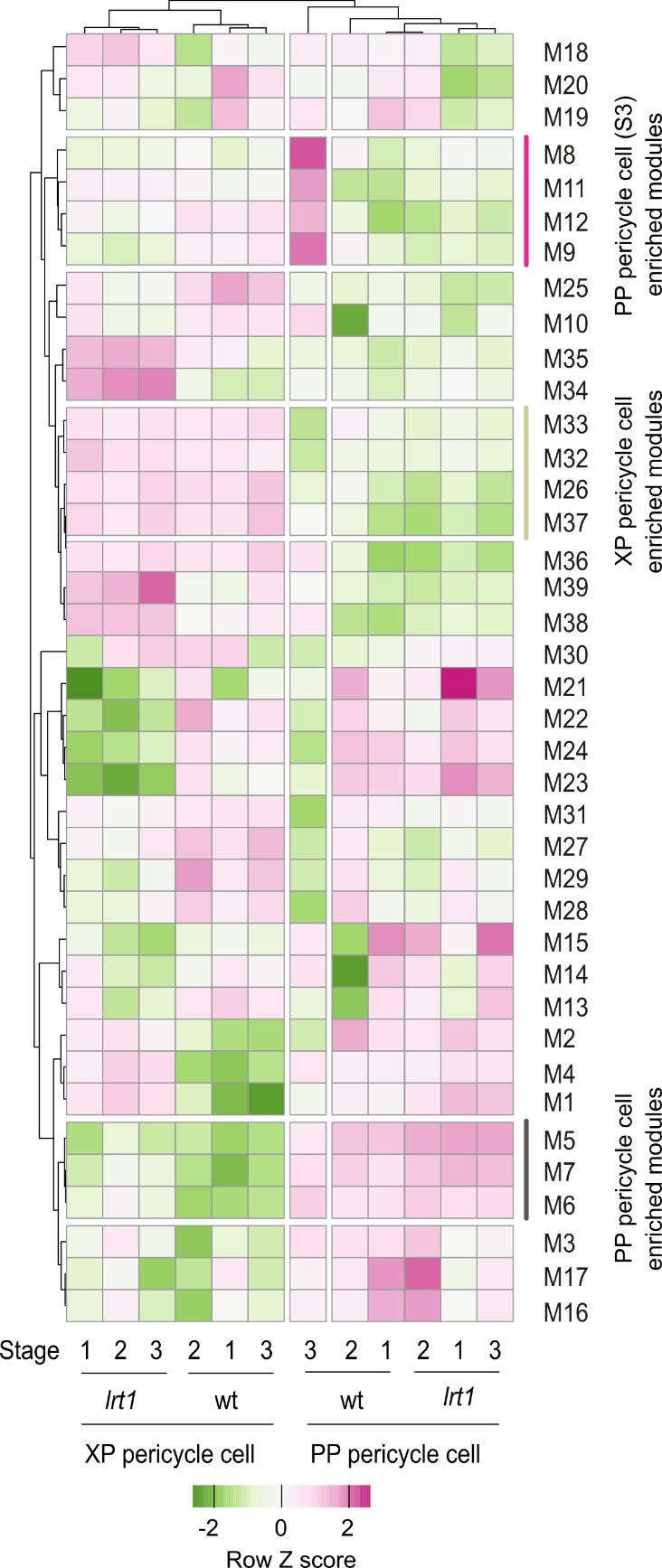

**Supplementary Figure** **1**. **Weighted gene co-expression network analysis (WGCNA) identifies stage- and genotype-specific transcriptional modules associated with pericycle cell competence.**
Heatmap showing correlations between module eigengenes (rows, M1–M39) and developmental traits (columns) across pericycle cell types (phloem-pole and xylem-pole), developmental stages (I–III), and genotypes (*lrt1* mutant and wild type). The color scale indicates Spearman correlation coefficients (SCC) ranging from negative (green) to positive (pink) correlations. Modules exhibiting strong positive correlations (SCC > 0.5) with specific pericycle cell types or developmental stages indicate transcriptional programs tightly associated with cell division competence and lateral root initiation. Notably, modules M8, M9, M11, and M12 are enriched in Stage III phloem-pole pericycle cells of wild-type roots, consistent with their transcriptional activation during pericycle cell transition into actively dividing states. These modules were further analyzed for transcription factor enrichment and motif composition, revealing a predominant role for the ERF gene family in regulating pericycle developmental reprogramming. PPPC, phloem-pole pericycle cell; XPPC, xylem-pole pericycle cell.

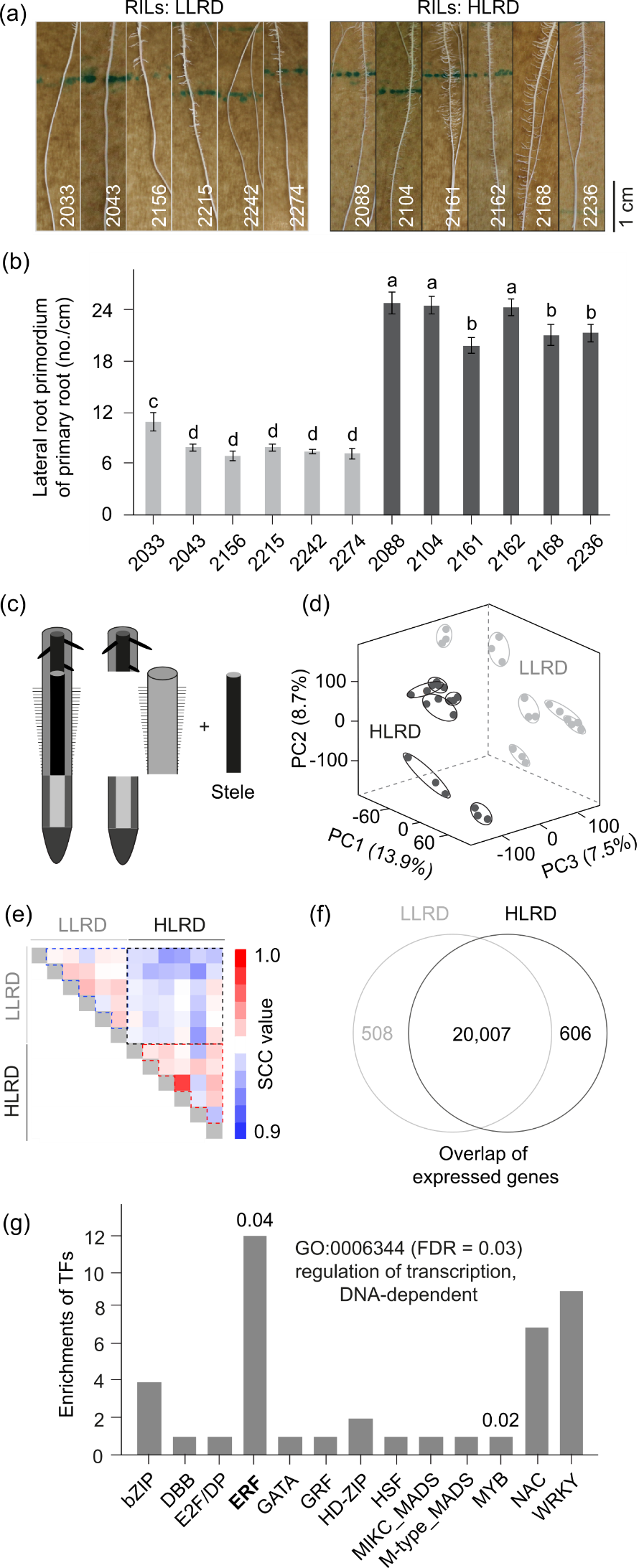

**Supplementary Figure** **2**. **Transcriptome profiling of stele tissue (comprising pecicycle) from recombinant inbred lines (RILs) differing in lateral root density reveals transcriptional signatures associated with root branching. (a)** RNA-seq was performed on manually dissected stele tissue from six maize RILs with low lateral root density and six with high lateral root density, selected from the IBM population. **(b)** Lateral root primordia phenotype of these six RILs. **(c)** Schematic illustration of root stele separation from non-stele tissue. **(d)** Principal component analysis (PCA) demonstrates a clear separation of transcriptomes according to lateral root phenotype, with PC1 explaining the majority of variation between groups. **(e)** Heatmap illustrates that spearman correlation coefficient (SCC) of gene expression among these RILs. **(f)** Venn diagram showing the number of genes exclusively expressed in high- and low-density RIL groups. A total of 606 genes were specifically expressed in high-density RILs, while 508 genes were enriched in low-density RILs. **(g)** GO enrichment analysis of high-density-specific genes highlights significant overrepresentation of transcriptional regulators, particularly ERF and MYB transcription factor families (Fisher’s exact test *p* < 0.05), suggesting their potential role in promoting lateral root initiation. FDR, false discovery rate.

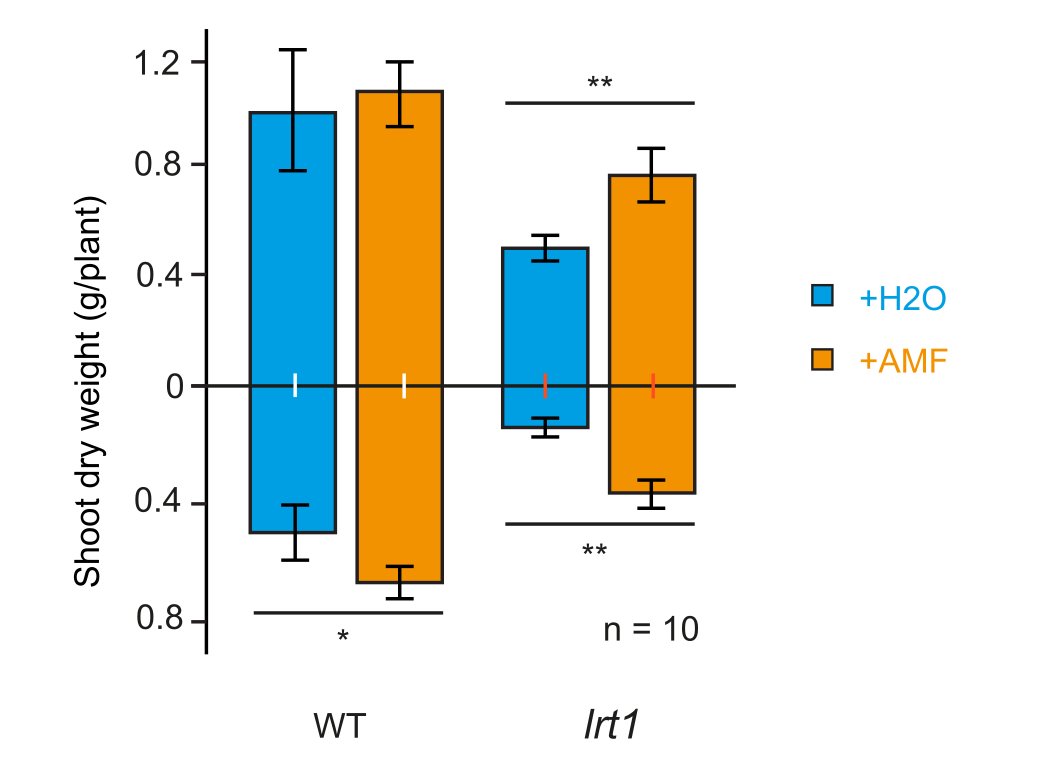

**Supplementary Figure** **3**. **Arbuscular mycorrhizal fungi (AMF) promote shoot biomass in maize *lrt1* mutants under phosphate-poor conditions.**

Shoot dry weight was measured in 6-week-old wild type (WT) and *lrt1* mutant plants grown in phosphate-poor artificial soil, with or without inoculation of arbuscular mycorrhizal fungi (AMF). AMF treatment significantly increased shoot dry biomass in both WT and *lrt1* plants compared to the water control (+H₂O). Notably, *lrt1* mutants showed a more pronounced biomass recovery upon AMF inoculation. Data represent mean ± SE (*n* = 10 plants per genotype per treatment). Statistical significance was assessed by two-sided Student’s *t*-test: *p* < 0.05 (*), *p* < 0.01 (**).

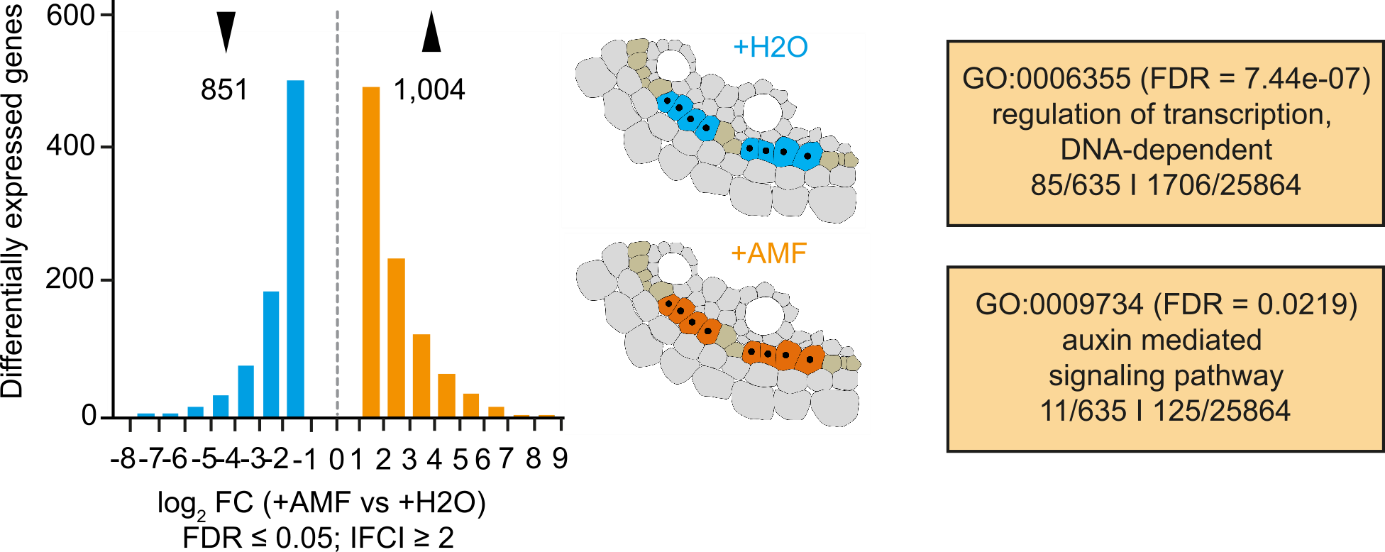

**Supplementary Figure** **4**. **AM fungi trigger transcriptomic reprogramming in *lrt1* pericycle cells, enriching transcriptional and auxin signaling pathways.**

Bar plots display the number of differentially expressed genes (DEGs) in *lrt1* pericycle cells upon AM fungal inoculation (+AMF, orange) compared to the water control (+H₂O, blue). A total of 1,004 genes were upregulated and 851 genes were downregulated (|FC| ≥ 2, FDR ≤ 0.05). Schematics indicate the sampled phloem-pole pericycle region captured by LCM for transcriptomic profiling. GO enrichment analysis of upregulated genes in response to AM fungi identified significant overrepresentation of genes involved in “regulation of transcription, DNA-dependent” (GO:0006355; FDR = 7.44e‑07) and “auxin-mediated signaling pathway” (GO:0009734; FDR = 0.0219), suggesting transcriptional reprogramming of hormone-responsive networks underlying AMF-induced lateral root development in *lrt1*.

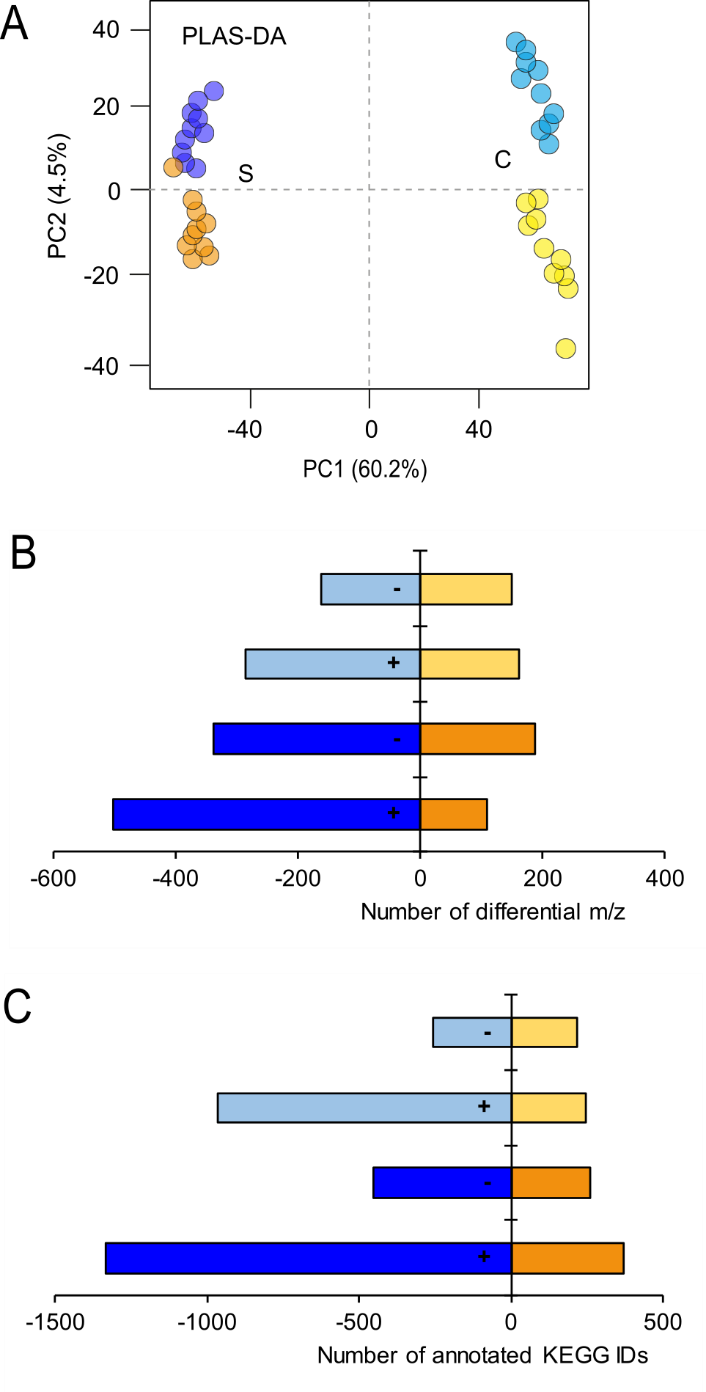

**Supplementary Figure** **5 Metabolomic reprogramming of *lrt1* roots in response to AM fungal colonization across tissues.**

(**A**) Principal coordinates analysis (PCoA) of metabolite profiles in *lrt1* root cortex and stele tissues inoculated with or without AM fungi. Each dot represents a distinct metabolic feature (m/z), colored by tissue and treatment. AMF inoculation induces distinct shifts in metabolic composition, particularly in cortex tissue.

(**B**) Bar plots showing the number of significantly differentially abundant mass features (m/z) in cortex (right groups) and stele (left groups) tissues, grouped by treatment (+/− AMF) and tissues (stele vs cortex). Positive (+) and negative (−) bars indicate upregulated or downregulated m/z features, respectively.

(**C**) Corresponding numbers of differentially annotated KEGG metabolic IDs based on the same comparisons shown in (B), highlighting AMF-driven reprogramming of known metabolic pathways, particularly in the cortex tissue.

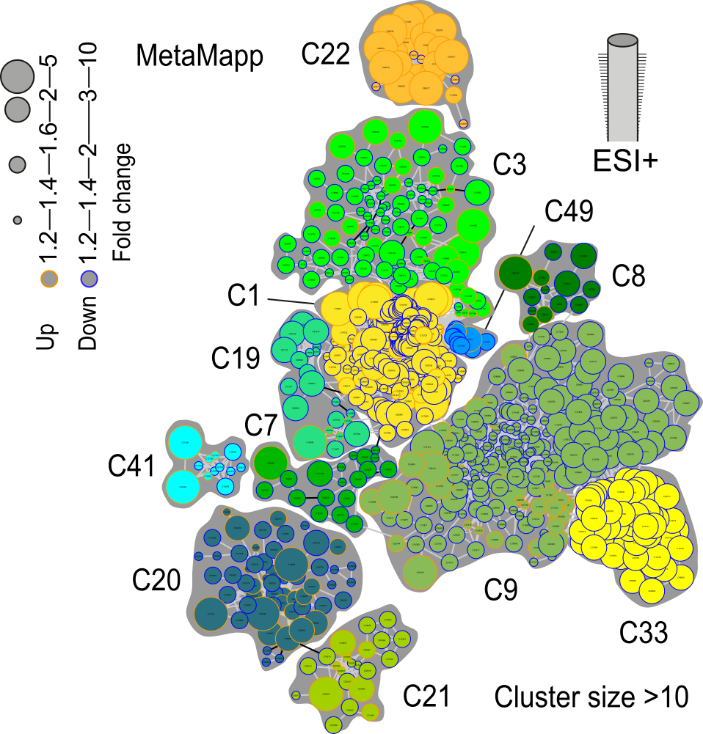

**Supplementary Figure** **6 Metabolic network of maize root cortex metabolites in response to AM fungal colonization.**

The network map was generated using MetaMapp based on chemical similarity and known biochemical relationships among detected metabolites. Each node represents a unique metabolite detected in the cortex tissue of *lrt1* mutant roots, and edges connect structurally or biochemically related metabolites. Node color denotes major chemical classes (e.g., flavonoids in blue/purple, sesquiterpenes in yellow, amino acid derivatives in green, fatty acids in cyan), while node size reflects the magnitude of fold change in abundance between AM fungi-inoculated and non-inoculated samples. Only metabolites significantly altered (VIP ≥1, FC ≥1.2 or ≤0.8, *q* < 0.05) are shown.

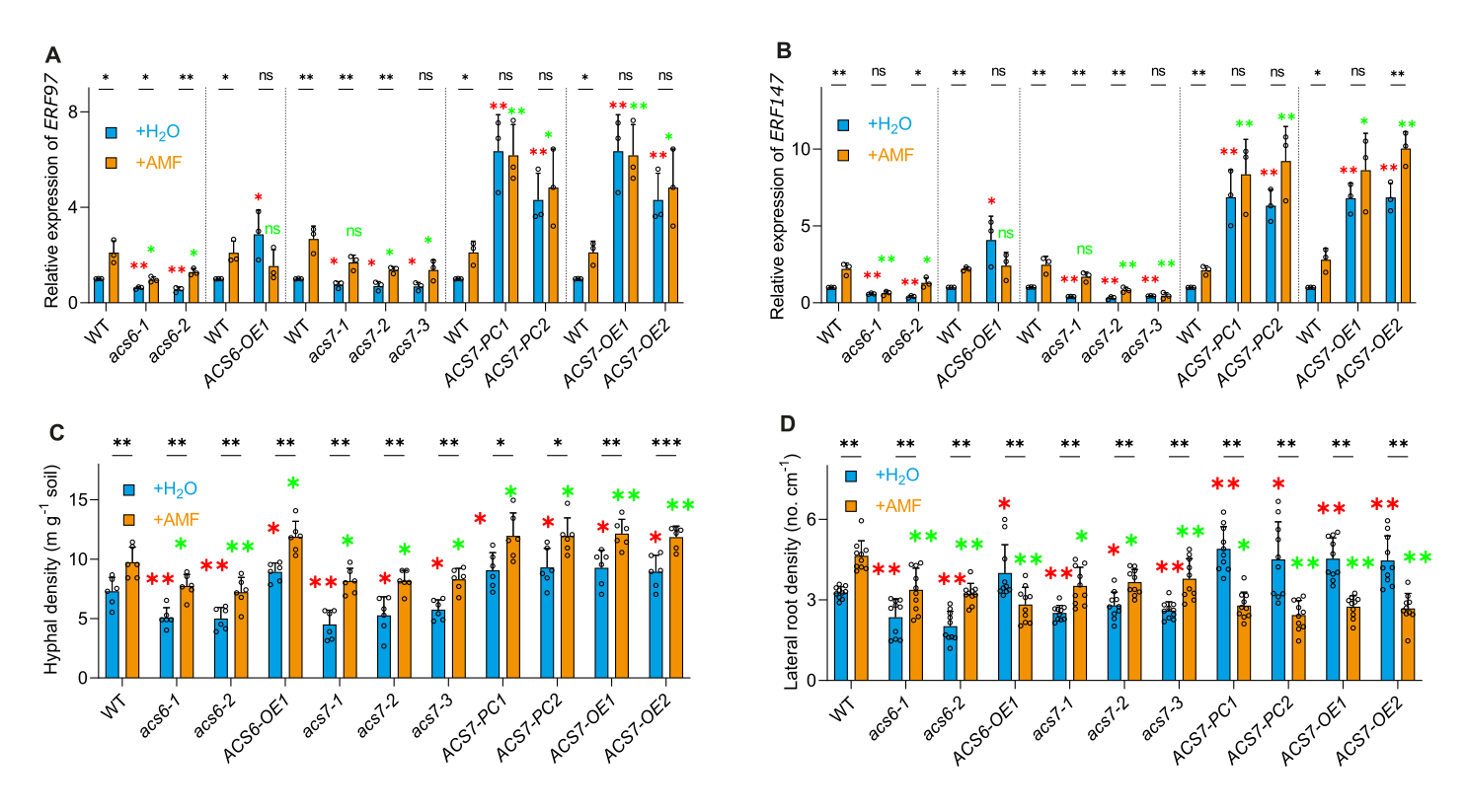
**Supplementary Figure** **7 Ethylene biosynthesis via ACS6/ACS7 modulates ERF activation, hyphal proliferation, and lateral root development under AMF colonization.**

(A, B) Expression levels of *ERF97* (A) and *ERF147* (B) quantified by qRT-PCR in roots of indicated genotypes, with or without AMF inoculation. Both ERF genes are significantly upregulated upon AMF colonization in wild type and overexpression lines, but this induction is impaired in *acs6-1* and *acs7-1* mutants.

(C) Hyphal length density (m g⁻¹ soil) in the rhizosphere. AMF-induced hyphal proliferation is significantly reduced in *acs6-1* and *acs7-1* mutants, but enhanced in overexpression and promoter construct lines, indicating ethylene biosynthesis promotes extraradical hyphal development.

(D) Lateral root density (no. cm⁻¹) is increased in WT and ACS-OE/PC lines upon AMF colonization, but not in *acs6-1* or *acs7-1* mutants, showing that ethylene is required for AM-induced root branching.

Bars represent mean ± SD (n = 4–6).
Student’s t-test was used for pairwise comparisons between treatments.
Significance is indicated as:
Green asterisks – AMF treatment effect (*P < 0.05, **P < 0.01),
Red asterisks – genotype effect,
ns – not significant.

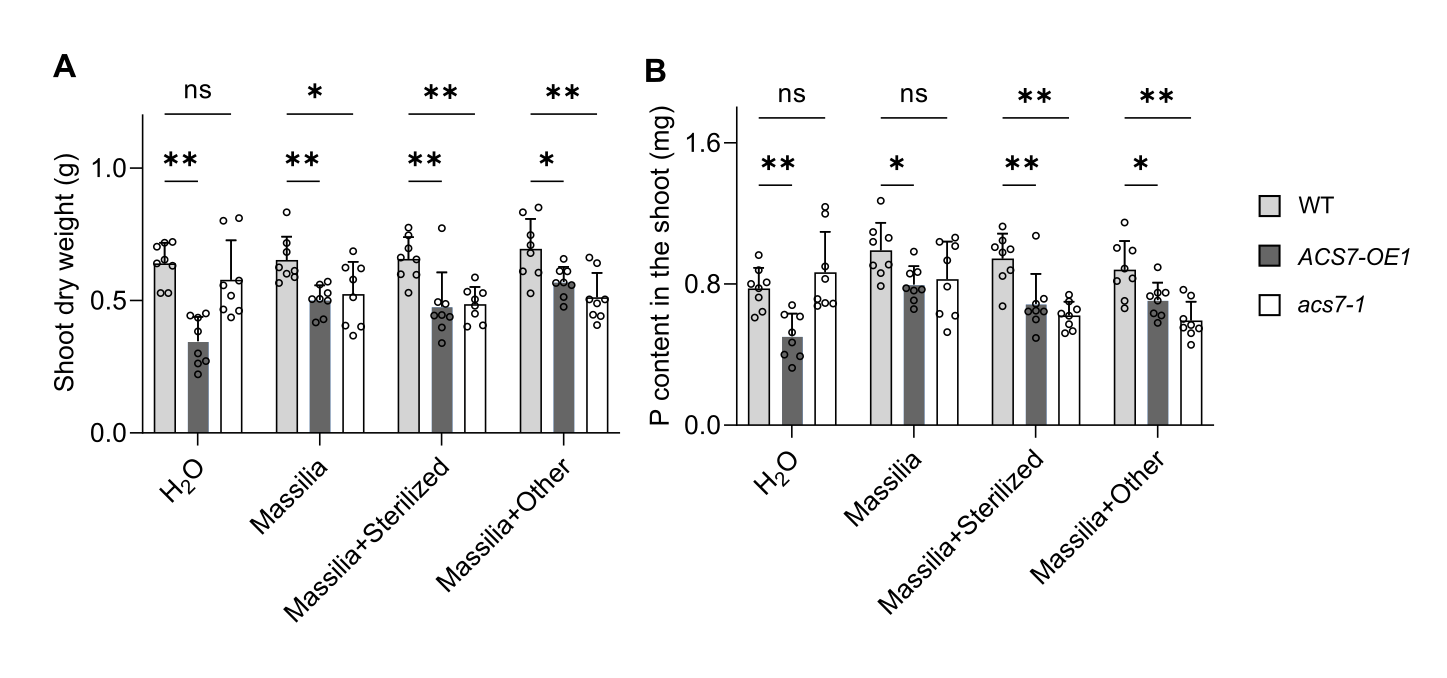

**Supplementary Figure** **8** Despite the negative growth and phosphorus acquisition phenotype of ACS7 overexpression, *Massilia* inoculation compensates by restoring both shoot biomass (A) and phosphorus uptake (B).

**Supplementary Table** **1**. Functional characterization of pericycle cell type specifically enriched modules (M5, M6, M7, M26, M32 and M33). PPPC, phloem-pole pericycle cell; XPPC, xylem-pole pericycle cell. FDR, false discovery rate.

| Module (M)  # | Cell type enriched | Module genes # | Enriched GO (FDR <0.05) | Enriched pathway | Gene ID | Gene name ^a^ |
| --- | --- | --- | --- | --- | --- | --- |
| M5 | PPPC | 1630 | GO: 0022402  (FDR = 0.045) | Cell cycle process | GRMZM2G128938 | atr1 |
|  |  |  |  |  | GRMZM2G037926 | cki2 |
|  |  |  |  |  | GRMZM2G157510 | cki3 |
|  |  |  |  |  | GRMZM2G158526 | cenH3 |
|  |  |  |  |  | GRMZM2G333916 | met1 |
|  |  |  |  |  | GRMZM5G822970 | mmus2 |
|  |  |  |  |  | GRMZM2G173162 | msh4 |
|  |  |  |  |  | GRMZM2G033828 | rrb3 |
|  |  |  |  |  | GRMZM2G129913 | spo1 |
|  |  |  |  |  | GRMZM2G157817 | xrcc3 |
|  |  |  |  |  | GRMZM2G395736 | - |
|  |  |  |  |  | GRMZM2G334857 | - |
|  |  |  |  |  | GRMZM2G134694 | - |
|  |  |  |  |  | GRMZM2G472023 | - |
|  |  |  |  |  | GRMZM2G100473 | - |
|  |  |  |  |  | GRMZM2G326328 | - |
|  |  |  |  |  | GRMZM2G300709 | - |
|  |  |  |  |  | GRMZM2G025340 | - |
|  |  |  |  |  | GRMZM2G133969 | - |
| M6 | PPPC | 275 | GO: 0015630  (FDR = 0.048) | Microtubule cytoskeleton | GRMZM2G436981 | krp11 |
|  |  |  |  |  | GRMZM2G020216 | mpk2 |
|  |  |  |  |  | GRMZM2G104704 | - |
|  |  |  |  |  | GRMZM2G334142 | - |
|  |  |  |  |  | GRMZM2G091543 | - |
|  |  |  |  |  | GRMZM2G471186 | - |
|  |  |  |  |  | GRMZM2G332809 | - |
|  |  |  |  |  | GRMZM2G555108 | - |
| M7 | PPPC | 1900 | GO: 0015630  (FDR = 0.00269) | Regulation of mitotic metaphase/anaphase transition | GRMZM5G899300 | bub3 |
|  |  |  |  |  | GRMZM2G047143 | mad2 |
|  |  |  |  |  | GRMZM2G022065 | - |
|  |  |  |  |  | GRMZM2G170591 | - |
|  |  |  |  |  | GRMZM2G009913 | - |
|  |  |  |  |  | GRMZM2G112072 | - |
| M26 | XPPC | 1,864 | GO: 0009832  (FDR = 0.014) | Plant-type cell wall biogenesis | GRMZM2G339122 | expa1 |
|  |  |  |  |  | GRMZM2G074585 | expa3 |
|  |  |  |  |  | GRMZM2G112336 | cesa1 |
|  |  |  |  |  | GRMZM2G027723 | cesa2 |
|  |  |  |  |  | GRMZM2G424832 | cesa4 |
|  |  |  |  |  | GRMZM2G070360 | dba2 |
|  |  |  |  |  | GRMZM2G073725 | gap1 |
|  |  |  |  |  | GRMZM2G013324 | gpx11 |
|  |  |  |  |  | GRMZM2G026218 | mmp187 |
|  |  |  |  |  | GRMZM2G069047 | nactf115 |
|  |  |  |  |  | GRMZM2G102946 | rop5 |
|  |  |  |  |  | GRMZM5G898668 | xt5 |
|  |  |  |  |  | GRMZM2G135743 | - |
|  |  |  |  |  | GRMZM5G804575 | - |
|  |  |  |  |  | GRMZM2G076394 | - |
|  |  |  |  |  | GRMZM2G007953 | - |
|  |  |  |  |  | GRMZM2G030167 | - |
|  |  |  |  |  | GRMZM2G001079 | - |
|  |  |  |  |  | GRMZM2G020721 | - |
|  |  |  |  |  | AC233901.1_FG008 | - |
| M32 | XPPC | 131 | GO: 0009755  (FDR = 0.024) | hormone-mediated signaling pathway | GRMZM2G104176 | iaa7 |
|  |  |  |  |  | GRMZM2G031615 | iaa14 |
|  |  |  |  |  | GRMZM2G180328 | [nactf20](https://maizegdb.org/data_center/locus/2775812) |
|  |  |  |  |  | GRMZM2G137046 | [bzip61](https://www.maizegdb.org/data_center/locus/2776589) |
|  |  |  |  |  | GRMZM2G042429 | - |
|  |  |  |  |  | GRMZM2G145518 | - |
| M33 | XPPC | 268 | GO: 0009734  (FDR = 0.014) | auxin mediated signaling pathway | GRMZM2G001799 | iaa41 |
|  |  |  |  |  | GRMZM5G825707 | iaa6 |
|  |  |  |  |  | GRMZM2G074267 | pin2 |
|  |  |  |  |  | GRMZM2G061515 | aas6 |
|  |  |  |  |  | GRMZM2G002515 | bri2 |
|  |  |  |  |  | GRMZM2G378580 | [arftf13](https://www.maizegdb.org/data_center/locus/2774535) |
|  |  |  |  |  | GRMZM2G012636 | - |

^a^ gene name in maize GDB.

**Supplementary Table 2.** MEME enriched motifs within 1 kb upstream and 0.5 kb downstream of promoter region of the pericycle cell division (stage 3) specifically enriched module (M9, M11, M12) genes.

| Module (M)  # | Motif # | E-value | Sites | Width | Sequence logo |
| --- | --- | --- | --- | --- | --- |
| M9 | 1 | 1.0e-089 | 97 | 45 | 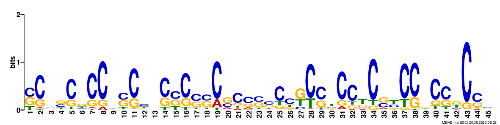 |
|  | 2 | 6.8e-061 | 96 | 29 | 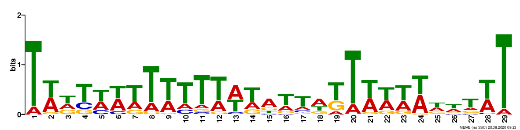 |
|  | 3 | 1.3e-077 | 86 | 50 | 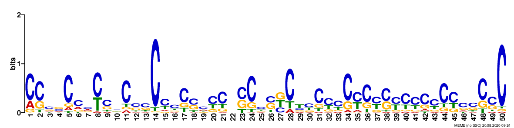 |
|  | 4 | 4.9e-043 | 62 | 29 | 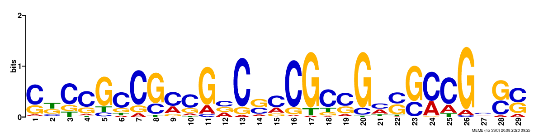 |
|  | 5 | 5.9e-035 | 75 | 29 | 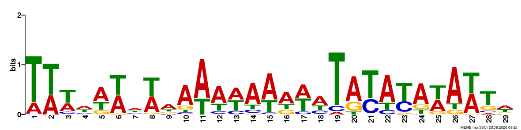 |
|  | 6 | 7.0e-028 | 88 | 20 | 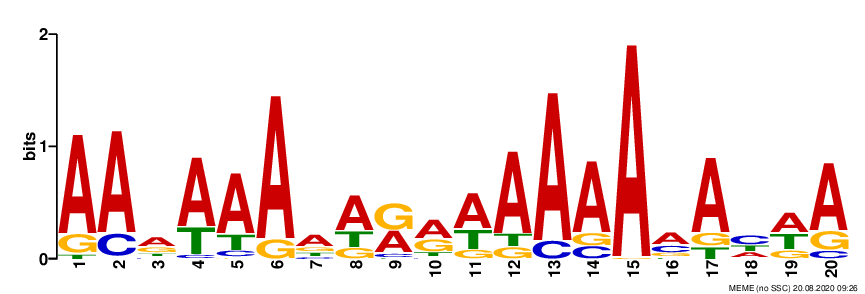 |
|  | 7 | 6.6e-016 | 102 | 21 | 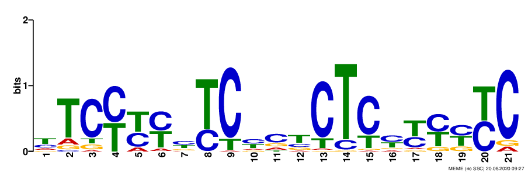 |
|  | 8 | 1.0e-018 | 93 | 28 | 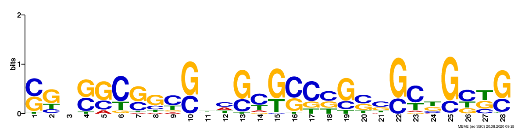 |
|  | 9 | 5.1e-010 | 106 | 29 | 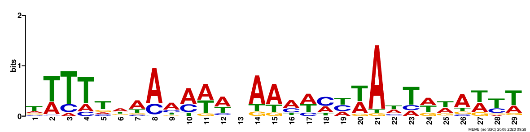 |
|  | 10 | 2.7e-008 | 57 | 29 | 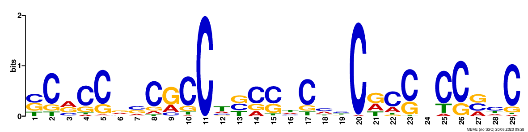 |
| M11 | 1 | 9.8e-084 | 79 | 29 | 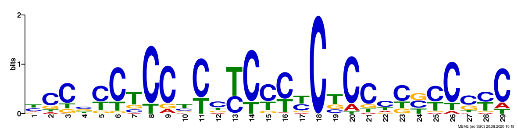 |
|  | 2 | 1.2e-081 | 91 | 41 | 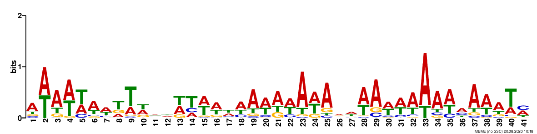 |
|  | 3 | 4.5e-080 | 83 | 29 | 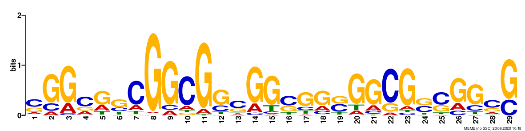 |
|  | 4 | 2.1e-033 | 58 | 29 | 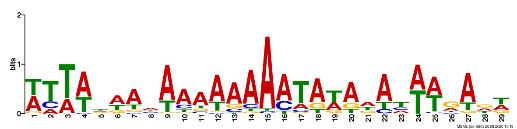 |
|  | 5 | 4.4e-026 | 61 | 21 | 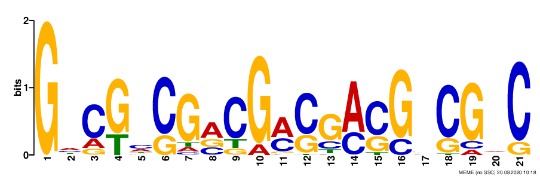 |
|  | 6 | 5.3e-039 | 88 | 37 | 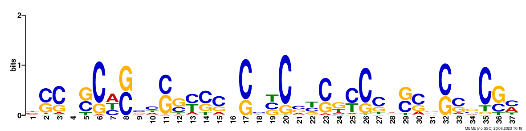 |
|  | 7 | 2.1e-015 | 56 | 15 | 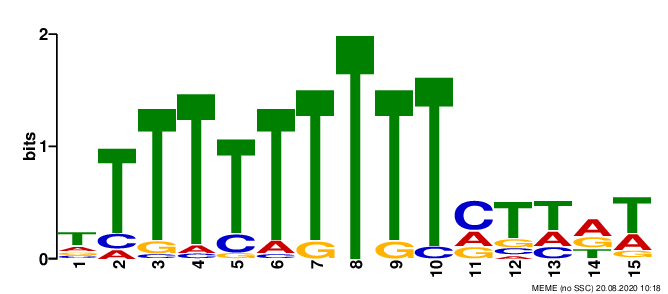 |
|  | 8 | 1.9e-014 | 60 | 29 | 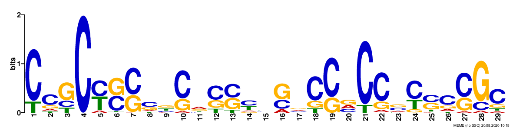 |
|  | 9 | 4.6e-007 | 68 | 21 | 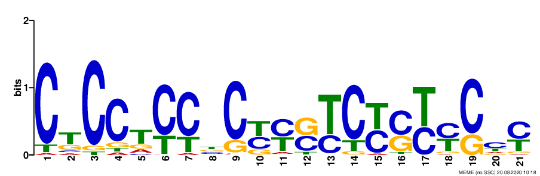 |
|  | 10 | 2.6e-006 | 28 | 21 | 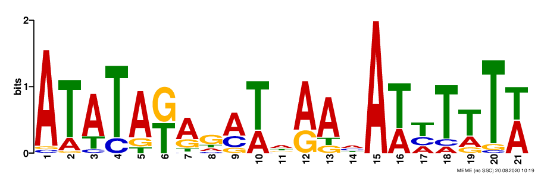 |
| M12 | 1 | 4.9e-104 | 143 | 29 | 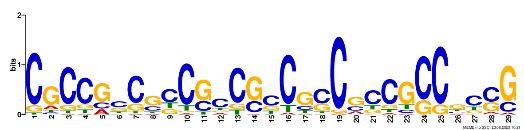 |
|  | 2 | 1.0e-098 | 107 | 29 | 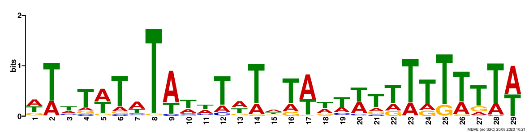 |
|  | 3 | 9.0e-067 | 87 | 41 | 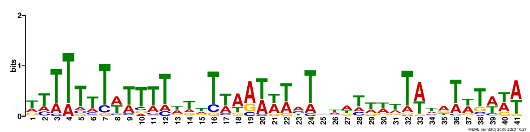 |
|  | 4 | 6.9e-046 | 157 | 21 | 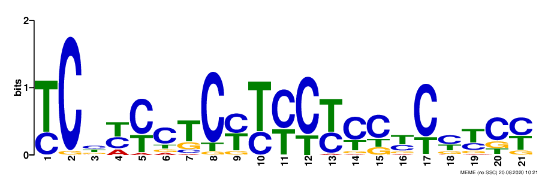 |
|  | 5 | 2.5e-040 | 149 | 37 | 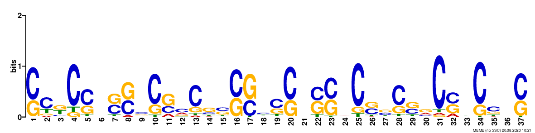 |
|  | 6 | 1.8e-036 | 121 | 21 | 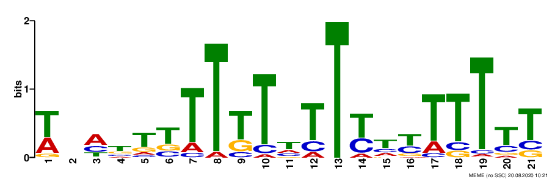 |
|  | 7 | 6.2e-020 | 155 | 21 | 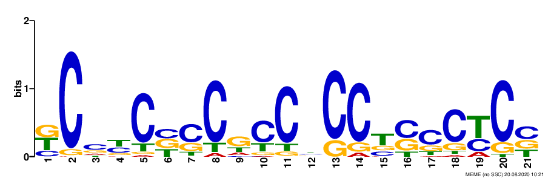 |
|  | 8 | 1.7e-011 | 111 | 29 | 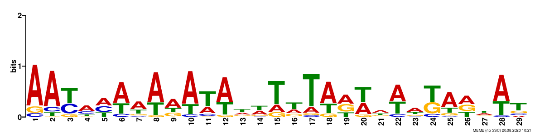 |
|  | 9 | 2.5e-028 | 70 | 41 | 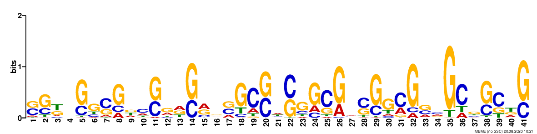 |
|  | 10 | 6.4e-006 | 23 | 21 | 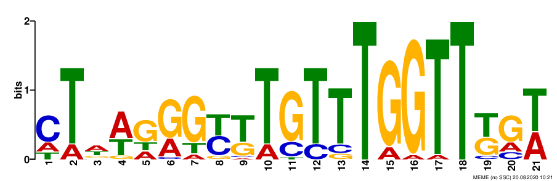 |

**Supplementary Table** **3**. TOMTOM identified top twenty *cis*-motifs with the most significant matches to the ERF transcription factors using JASPAR CORE Plants database. TF, transcription factor.

| Module (M)  # | Input Motif # | Total matches | Significant matches (*q*-value< 0.05) | AP2/ERF domain Class | Matching TF^a^ | *q*-value |
| --- | --- | --- | --- | --- | --- | --- |
| M9 | 1 | 55 | 38 | 36 | **ERF104** | 2.44e-03 |
|  | 4 | 64 | 54 | 52 | ERF9 | 4.33e-07 |
|  | 8 | 48 | 32 | 32 | ERF3 | 1.06e-02 |
|  | 10 | 71 | 51 | 50 | ERF4 | 8.05e-04 |
| M11 | 3 | 61 | 54 | 54 | **ERF9** | 1.25e-07 |
|  | 5 | 56 | 27 | 26 | ERF104 | 5.22e-3 |
|  | 6 | 48 | 11 | 10 | CRF4 | 8.16e-03 |
|  | 8 | 52 | 4 | 4 | ERF4 | 2.97e-02 |
| M12 | 1 | 59 | 54 | 54 | **ERF9** | 2.77e-08 |
|  | 5 | 64 | 44 | 41 | CRF2 | 7.39e-04 |
|  | 7 | 59 | 30 | 30 | ERF104 | 9.40e-03 |
|  | 9 | 63 | 46 | 46 | ERF104 | 2.02e-3 |

^a^ Transcription factors of which the known binding motif most significantly matches the identified motif (*q*-value < 0.05) in Arabidopsis.

**Supplementary Table** **4**. Functional characterization of proteins exhibiting stele-specific enrichment in response to AM fungi inoculation. “+” enriched in response to AM fungi; “-” depleted in response to AM fungi.

| Protein ID | KEGG | Description | GO (Biological Process) | Enrichment |
| --- | --- | --- | --- | --- |
| tr\|A0A1D6NEY9\|A0A1D6NEY9_MAIZE | K00799 | glutathione S-transferase | GO:0048527, lateral root development | 0.74 (-) |
| tr\|K7TXJ3\|K7TXJ3_MAIZE | K01807 | ribose 5-phosphate isomerase A | GO:0055085, transmembrane transport | 0.12 (-) |
| tr\|Q9M588\|Q9M588_MAIZE | K17080 | prohibitin 1 | GO:0048527, lateral root development | 0.8 (-) |
| tr\|B4FIE9\|B4FIE9_MAIZE | K00789 | S-adenosylmethionine synthetase | GO:0006556, S-adenosylmethionine biosynthetic process | 1.3 (+) |
| tr\|B6SIK2\|B6SIK2_MAIZE | / | / | / | 1.51 (+) |
| tr\|B4FVJ0\|B4FVJ0_MAIZE | K02930 | large subunit ribosomal protein L4e | GO:0006412, translation | 1.24 (+) |
| tr\|B6SJ21\|B6SJ21_MAIZE | K14753 | guanine nucleotide-binding protein subunit beta-2-like 1 protein | GO:0048364, root development | 1.28 (+) |

**Supplementary Table** **5**. Proteins exhibiting enrichment in cortex tissue in response to AM fungi inoculation are annotated to Phenylpropanoid biosynthesis. “+” enriched in response to AM fungi; “-” depleted in response to AM fungi.

| **Protein ID** | **Description** | **Enrichment** | **Coding gene ID** |
| --- | --- | --- | --- |
| tr\|B6TBU1\|B6TBU1_MAIZE | 1-aminocyclopropane-1-carboxylate oxidase | 1.52 (+) | AC148152.3_FG005 |
| sp\|P49235\|HGGL1_MAIZE | 4-hydroxy-7-methoxy-3-oxo-3,4-dihydro-2H-1,4-benzoxazin-2-yl glucoside beta-D-glucosidase 1, chloroplastic | 1.41 (+) | GRMZM2G016890 |
| tr\|A0A1D6EZ64\|A0A1D6EZ64_MAIZE | Polygalacturonase inhibitor 1 | 1.47 (+) | GRMZM2G099295 |
| tr\|A0A1D6HY96\|A0A1D6HY96_MAIZE | Serine carboxypeptidase-like 19 | 1.4 (+) | GRMZM2G471269 |
| tr\|A0A1D6MUP9\|A0A1D6MUP9_MAIZE | Beta-glucosidase 17 | 1.34 (+) | GRMZM2G426467 |
| tr\|A0A1D6IXD5\|A0A1D6IXD5_MAIZE | Beta-glucosidase2 | 1.43 (+) | GRMZM2G008247 |
| tr\|A0A1D6GWL6\|A0A1D6GWL6_MAIZE | GDSL esterase/lipase | 1.39 (+) | GRMZM2G132331 |
| tr\|B4FHG3\|B4FHG3_MAIZE | Peroxidase | 1.33 (+) | GRMZM2G108219 |
| tr\|A0A1D6E530\|A0A1D6E530_MAIZE | Peroxidase | 1.34 (+) | GRMZM2G108153 |
| tr\|C0HHA6\|C0HHA6_MAIZE | Peroxidase | 1.34 (+) | GRMZM2G122853 |
| tr\|B6T3V1\|B6T3V1_MAIZE | Peroxidase | 1.37 (+) | GRMZM2G089982 |
| tr\|C4J6E4\|C4J6E4_MAIZE | Peroxidase | 1.39 (+) | GRMZM2G126261 |
| tr\|B6SIU4\|B6SIU4_MAIZE | Peroxidase | 1.43 (+) | GRMZM2G144648 |
| tr\|A0A1D6F4C8\|A0A1D6F4C8_MAIZE | Peroxidase | 1.46 (+) | GRMZM2G133475 |
| tr\|A0A1D6LYW3\|A0A1D6LYW3_MAIZE | Peroxidase | 1.56 (+) | GRMZM2G135108 |
| tr\|D7NLB3\|D7NLB3_MAIZE | Peroxidase | 1.62 (+) | GRMZM2G410175 |
| tr\|B4FG39\|B4FG39_MAIZE | Peroxidase | 1.67 (+) | GRMZM2G108123 |
| tr\|A0A1D6KAW3\|A0A1D6KAW3_MAIZE | Peroxidase | 1.73 (+) | GRMZM2G088765 |
| sp\|Q9FEQ8\|PER2_MAIZE | Peroxidase 2 | 1.71 (+) | GRMZM2G040638 |
| tr\|A0A1D6HDL5\|A0A1D6HDL5_MAIZE | Phenylalanine ammonia-lyase | 0.74 (-) | GRMZM2G029048 |

**Supplementary Table** **6**. Primers used for gene cloning, expression, and vector construction

| **Gene** | **Primer (5' to 3')** | **Accession #** |
| --- | --- | --- |
| *ZmACS6_qF* | GGCAGCATCATTAGCAGTAGGA | *GRMZM2G054361/Zm00001d033862* |
| *ZmACS6_qR* | ACCACTCCTGCTCCTACCTAAT |  |
| *ZmACS6-CRISPR-F* | TGGGGTGGGAGGAGTATGA |  |
| *ZmACS6-CRISPR-R* | GTATGGCGTGGGGATGAGA |  |
| *ZmACS7_qF* | CGTACGTAGCTAGCTTGACATT | *GRMZM5G894619/Zm00001d026060* |
| *ZmACS7_qR* | TTGACTAGCTGCTTACCGCC |  |
| *ZmACS7-CRISPR-F* | AGCAAGCTCATGCCCTTCAA |  |
| *ZmACS7-CRISPR--R* | AGCATTGGACTTGGATCGGC |  |
| *ZmACS7-PC-F* | AGGGTGAAGCTCAACGTGTC |  |
| *ZmACS7-PC-R* | TGTCACTCTGTGTAATGTCAAGC |  |
| *ZmERF1_qF* | GGCTTCGAACTGGATCTGAACT | *GRMZM2G310368/Zm00001d043205* |
| *ZmERF1_qR* | GCCATACAGGACCAAAACAACC |  |
| *ZmERF2_qF* | TCGATCGTTTAATTTGCCTGCC | *GRMZM2G068967/Zm00001d002364* |
| *ZmERF2_qR* | TGTTTATACCACGTAGCCTGCC |  |
| *C2_qF* | CTGGCCTCATCTCCAAGAACAT | *GRMZM2G422750/Zm00001d052673* |
| *C2_qR* | CATCTCGTCGAGGATGAAGAGG |  |
| *qZmTublinF* | GCTATCCTGTGATCTGCCCTGA | *GRMZM2G152466/Zm00001d013367* |
| *qZmTublinR* | CGCCAAACTTAATAACCCAGTA |  |
| *F1048* | GTGSTGCAYGGYTGTCGTCA |  |
| *R1194* | ACGTCRTCCMCACCTTCCTC |  |
| *qRT_Massilia_FW* | GGAGCGGCCGATRTCTGATTAG |  |
| *qRT_Massilia_RV* | ATTCTACCCCCCTCTGCCARAY |  |
